# Lipopolysaccharide chain length and coat assembly keep guanylate-binding proteins upright on bacterial membranes

**DOI:** 10.64898/2026.10.29.762752

**Authors:** Bastian Ferdinand Bundschuh, Jennifer Loschwitz, Birgit Strodel

## Abstract

Guanylate-binding proteins (GBPs) defend host cells by assembling on intracellular pathogen membranes. For human GBP1, this requires a switch from a compact, closed conformation to an extended, open one, whereas for murine orthologues the open state is not known. How the open state behaves at a membrane has remained inaccessible to experiment. Using all-atom and coarse-grained molecular dynamics simulations covering over 0.5 ms, we followed human GBP1 and murine GBP2 as monomers, dimers, and higher-order assemblies on phosphatidylcholine membranes, *Shigella flexneri* lipopolysaccharide (LPS), and phosphoinositide-containing bilayers. Closed GBPs dock through their prenyl anchor with minimal lipid selectivity. Open GBPs, by contrast, are intrinsically unstable: monomers fold back into membranebound resting states, and dimers tip over unless supported, either by long LPS O-antigen chains or by neighboring molecules within a coat. The open state thus appears to be maintained not by the nucleotide alone but by the protein’s molecular environment.

## 1 Introduction

Guanylate-binding proteins (GBPs) are large GTPases of the dynamin superfamily, induced by interferon-*γ* (IFN-*γ*) and central to the cell-autonomous immune response against intracellular pathogens including bacteria, viruses, and protozoa.^1^ Unlike most GTPases, which convert GTP to GDP, several GBPs further hydrolyze GDP to GMP via a sequential twostep mechanism, distinguishing them from classical small GTPases and underpinning their high intrinsic turnover rates.^2^ Structurally, GBPs comprise a GTPase domain (GD), a middle domain (MiD), and a helical effector domain (ED), with C-terminal CaaX motifs subject to isoprenylation — most commonly farnesylation or geranylgeranylation — that regulate membrane targeting.^3–8^ GTP binding and hydrolysis drive conformational changes that promote oligomerization and membrane association, enabling the formation of higher-order assemblies at pathogen-related membranes.^9^ Once recruited to intracellular pathogens or their vacuoles, GBPs cooperate with immunity-related GTPases (IRGs) to destabilize pathogen-containing vacuoles (PCVs), promote autophagy via ubiquitination and adaptors such as p62 and LC3, and recruit caspase-4 to initiate non-canonical inflammasome assembly.^10–14^

Among the seven human and eleven murine GBP paralogs, human GBP1 (hGBP1) and murine GBP2 (mGBP2) stand out as the best-characterized representatives of their respective species. They share 68% sequence identity (82% similarity) and the same two-step hydrolysis of GTP via GDP to GMP, although mGBP2 converts GDP to GMP less efficiently than hGBP1.^15,16^ Both carry a C-terminal CaaX motif — farnesylated in hGBP1 and geranylgeranylated in mGBP2 — and function as molecular hubs orchestrating broad antimicrobial programs against bacteria, parasites, and viruses.^16–18^ Despite their overall similarity, key differences exist: hGBP1 carries C-terminal polybasic segments enriched in lysines and arginines that tune pathogen targeting, whereas mGBP2 lacks this triple-arginine sequence and relies on a single lysine residue (K585) preceding the CaaX motif for membrane association.^17,19^ Sequence divergence is greatest in the C-terminal ED, which likely underlies these functional differences.^19,20^

A critical and incompletely understood aspect of GBP function is the transition between closed and open conformational states. The closed state, favored in the absence of nucleotide or upon GDP binding after hydrolysis, maintains a compact conformation that prevents oligomerization and membrane association, whereas nucleotide binding and hydrolysis induce an open state in which exposure of the ED drives dimerization and membrane attachment, enhancing antimicrobial activity.^16,21–27^ Previous molecular dynamics (MD) simulations of hGBP1 with bound GTP have shown that open monomers are stabilized by the nucleotide but can spontaneously revert to a closed state stabilized by interdomain salt bridges.^28^ Dimerization substantially increases the conformational stability of the open state through a criss-cross arrangement, in which the stalks (MiD and ED) of the two protomers cross each other at the GD–MiD transition, and through coiled-coil formation in the EDs.^23,28^ In larger hGBP1 assemblies, dimers serve as the building blocks, which are thought to interact via contacts between helix *α*4*^′^* and the neighboring GD. This interface is proposed to form the polymeric contacts that enable coat formation on pathogen membranes.^22,29^

While open states of hGBP1 have been proposed and partially characterized,^22,23,30^ open states of mGBP2 have not been demonstrated. Given the high sequence identity between the two proteins, we hypothesize that mGBP2 also adopts an open conformation, which is supported by the extended dimer conformation that AlphaFold 3^31^ predicts for mGBP2 and the related mGBP7, albeit without the criss-cross arrangement of the MiDs (**Fig. S1**). A second unresolved question concerns the membrane specificity of these proteins: whereas lipopolysaccharide (LPS) is the primary pathogen-associated molecular pattern (PAMP) recruiting hGBP1, the stimuli and binding mechanisms governing mGBP2 membrane association remain poorly understood. More broadly, how closed and open GBP monomers, dimers, and higher-order oligomers interact with pathogen-related membranes, and how they modulate membrane dynamics, has not yet been characterized at the molecular level.

To address these questions, we employ all-atom molecular dynamics (AAMD) and coarsegrained molecular dynamics (CGMD) simulations to investigate hGBP1 and mGBP2 across monomeric, dimeric, and oligomeric states at membranes of increasing complexity, from a pure 1-palmitoyl-2-oleoyl-*sn*-glycero-3-phosphocholine (POPC) reference to a realistic *Shigella flexneri* LPS membrane and an asymmetric membrane containing phosphatidylinositol 5-phosphate (PI5P). We characterize the binding positions and orientations of closed and open protein forms, assess how oligomeric assemblies, including hexamers, a periodic coat-like assembly, and a disc of 30 dimers modeled after the polymeric hGBP1 assembly resolved by cryo-electron tomography,^22^ interact with and affect the local properties of membranes, and compare behavior across membrane compositions. This work provides a molecular-level view of GBP–membrane interaction mechanisms that have so far been inferred mainly from experiments. The protein structures and membrane models used are introduced in **Fig. 1**.

**Figure 1:**
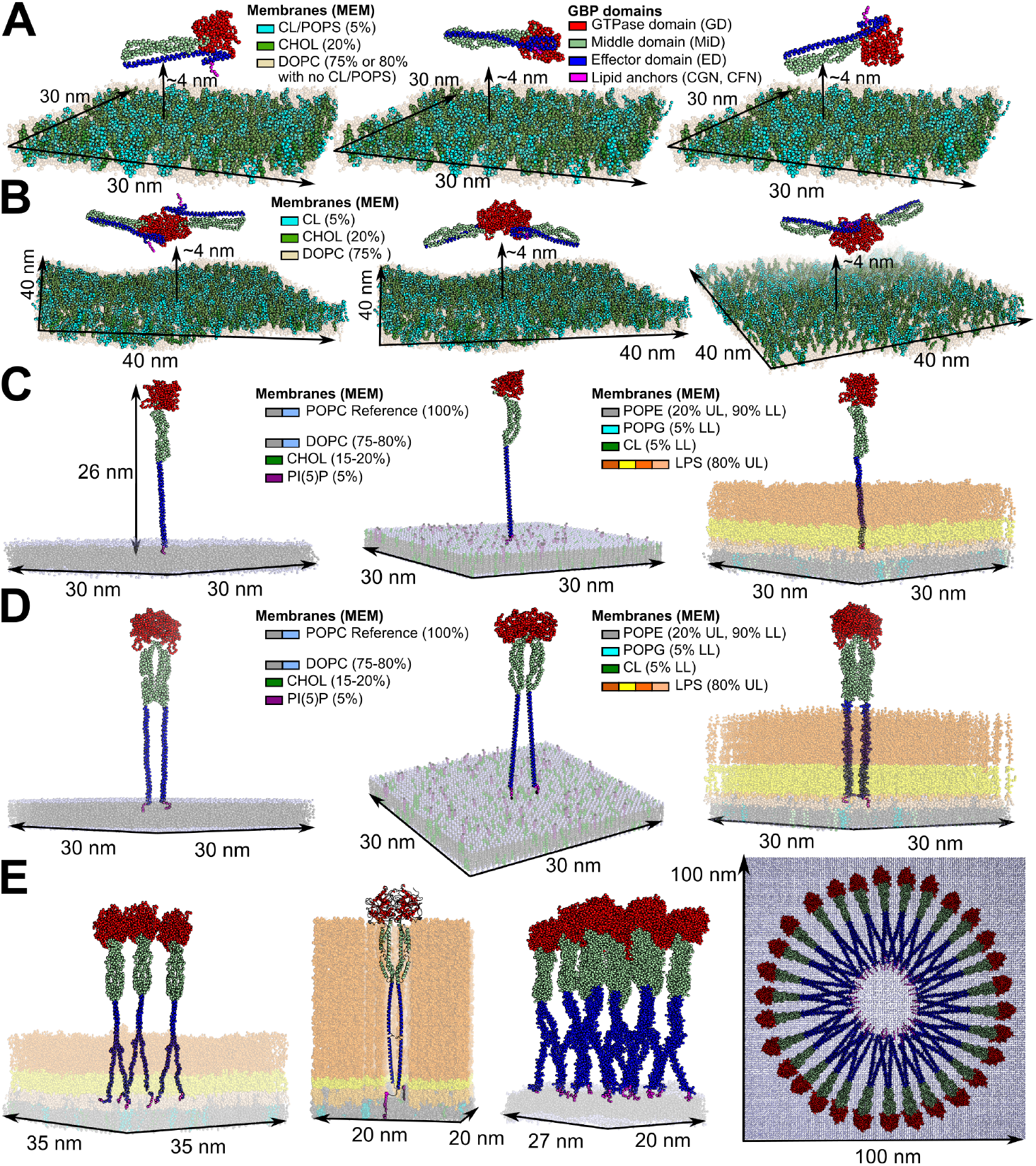
Systems for MD simulations. **(A)** The closed mGBP2 monomer systems in three different orientations on 30 *×* 30 nm^2^ membranes, where the mGBP2 was placed 4 nm away from the membrane. **(B)** The mGBP2 dimer systems show three different orientations on a 40 *×* 40 nm^2^ membrane with cardiolipin. **(C)** The open monomer systems of hGBP1 and mGBP2 on the 30 *×* 30 nm^2^ membranes, where the GBPs were inserted in the membrane with their prenylation. We used a pure POPC membrane as reference for both species and more realistic membranes depending on the organism. For hGBPs asymmetric LPS membranes were set up, i.e. upper leaflet with 80% LPS & 20% POPE; lower leaflet with 90% POPE, 5% POPG & 5% cardiolipin (CL). For the asymmetric PI5P membranes simulated together with mGBP2: 5% PI5P, 20% CHOL, 75% DOPC in the upper leaflet and 20% CHOL and 80% DOPC in the lower leaflet were set up. **(D)** The open dimer systems of hGBP1 and mGBP2 embedded in the same type of membranes as for the open monomer systems. **(E)** The open hexamer and dimer hGBP1 on LPS membranes with different O-antigen chain lengths. **(F)** The coat-like and disc-like systems of hGBP1 on pure POPC membranes. Membrane lipid compositions and GBP domains are given as legends in the figure and chemical identities of lipids can be found in (**Fig. S2**).

## 2 Results

All simulations were based on structural models of prenylated hGBP1 and mGBP2 in closed and open conformations. Closed-state coarse-grained (CG) models of mGBP2 were built from our previous studies^32,33^ using *martinize.py* and *insane.py*, while open-state structures of hGBP1 were taken from prior simulations^28^ and cryo-EM/cryo-ET structures (PDB: 8CQB, 8R1A);^22,23^ open-state models of mGBP2 were generated with AlphaFold 3^31^ and SwissModel,^34^ with extended domains and prenyl groups added in PyMOL.^35^

Membrane systems were constructed using CHARMM-GUI,^36^ covering a range of compositions from pure POPC to realistic *S. flexneri* LPS and asymmetric phosphatidylinositol 5-phosphate (PI5P)-containing bilayers. To be specific, the following membrane models were simulated: (i) a pure POPC membrane as reference, (ii) a membrane model composed of DOPC (1,2-dioleoyl-*sn*-glycero-3-phosphocholine), cholesterol (CHOL), and cardiolipin (CL) in a ratio of 75%/20%/5% (CL-mem), (iii) a model with DOPC, CHOL, and POPS (1-palmitoyl-2-oleoyl-*sn*-glycero-3-phospho-L-serine) in a ratio of 75%/20%/5% as a negative control for electrostatic effects of anionic lipids (PS-mem), (iv) a DOPC/-CHOL membrane in a ratio of 80%/20% as a reference (PC-mem), (v) a realistic *S. flexneri* LPS membrane containing 80% LPS and 20% POPE (1-palmitoyl-2-oleoyl-*sn*-glycero-3-phosphoethanolamine) in the upper leaflet and 90% POPE, 5% POPG (1-palmitoyl-2-oleoyl-*sn*-glycero-3-phosphoglycerol), and 5% CL in the lower leaflet, and (vi) an asymmetric PI5P-containing membrane model with 5% PI5P, 20% CHOL, and 75% DOPC in the upper leaflet and 20% CHOL and 80% DOPC in the lower leaflet. The different forms of mGBP2 or hGBP1 — encompassing closed and open conformations as well as monomeric, dimeric, and higher-order oligomeric states — were placed above these membrane models as shown in **Fig. 1**. The closed mGBP2 membrane systems, all systems containing LPS (the LPS reference membranes, the open hGBP1 monomers and dimers on LPS, and the hGBP1 hexamer on LPS), and, for direct comparison, the open hGBP1 monomers and dimers on POPC used the MARTINI 2.2 force field,^37,38^ as LPS is available only in this version. Open mGBP2 monomers and dimers and the larger hGBP1 assemblies in solution and on POPC (hexamer, 18mer, coat, and disc) employed MARTINI 3,^39^ to take advantage of updated phosphoinositide and prenylation parameters^40,41^ and improved protein–protein interactions. All-atom simulations using CHARMM36m^42^ were performed only for the open hGBP1 dimer on long O-antigen LPS membranes, where full-length LPS reaches the GD — a regime not yet accessible at the CG level. All systems underwent energy minimization and equilibration at 310 K and 1 bar, followed by production runs of 1 *µ*s (AAMD, 2-fs time step) or 1–10 *µ*s (CGMD, 20-fs time step), as listed per system in **Table 1**. Full methodological details are provided in the Methods.

**Table 1:** Summary of the AAMD and CGMD simulation systems performed in this study, organized by section.

| **Section 2.1 Binding of closed mGBP2 to lipid membranes** (Simulated with MARTINI 2.2) |  |  |  |  |
| --- | --- | --- | --- | --- |
| MD | System | size [atoms/beads] | run x length | cumulated time [μs] |
| CGMD | Membrane references | up to 200,000 | $3 \times 10 \mu$s | 30 μs |
| CGMD | mGBP2 monomer on membrane | $\sim 224,000$ | $27 \times 10 \mu$s | 270.0 μs |
| CGMD | mGBP2 dimer on membrane | $\sim 540,000$ | $3 \times 10 \mu$s | 30.0 μs |
| **Section 2.2 The open states in solution** (Simulated with MARTINI 3) |  |  |  |  |
| MD | System | size [atoms/beads] | run x length | cumulated time [μs] |
| CGMD | mGBP2 monomer/dimer in solution | $\sim 350,000$ | $2 \times 5 \mu$s | 10.0 μs |
| CGMD | mGBP7 monomer/dimer in solution | $\sim 600,000$ | $2 \times 5 \mu$s | 10.0 μs |
| CGMD | hGBP1 hexamer in solution | $\sim 350,000$ | $1 \times 5 \mu$s | 5.0 μs |
| **Section 2.3 The open states at membranes** (Simulated with MARTINI 2.2, MARTINI 3 and C36m) |  |  |  |  |
| MD | System | size [atoms/beads] | run x length | cumulated time [μs] |
| CGMD | Membrane references | up to 200,000 | $5 \times 10 \mu$s | 50 μs |
| CGMD | hGBP1/mGBP2 monomers on simple membrane | $\sim 750,000$ | $2 \times 10 \mu$s | 20.0 μs |
| CGMD | hGBP1/mGBP2 dimer on simple membrane | $\sim 750,000$ | $2 \times 10 \mu$s | 20.0 μs |
| CGMD | hGBP1/mGBP2 monomer on realistic membrane | $\sim 750,000$ | $2 \times 10 \mu$s | 20.0 μs |
| CGMD | hGBP1/mGBP2 dimer on realistic membrane | $\sim 750,000$ | $2 \times 10 \mu$s | 20.0 μs |
| AAMD | hGBP1 dimer on realistic membrane (long LPS) | $\sim 2,150,000$ | $2 \times 1 \mu$s | 2.0 μs |
| CGMD | hGBP1 hexamer on realistic membrane | $\sim 750,000$ | $1 \times 10 \mu$s | 10.0 μs |
| **Section 2.4 GBP polymerization** (Simulated with MARTINI 3) |  |  |  |  |
| MD | System | size [atoms/beads] | run x length | cumulated time [μs] |
| CGMD | hGBP1 hexamer/octadecamer on membrane | $\sim 2,500,000$ | $2 \times 5 \mu$s | 10.0 μs |
| CGMD | hGBP1 coat-like assembly on membrane | $\sim 1,500,000$ | $1 \times 10 \mu$s | 10.0 μs |
| CGMD | hGBP1 disc on membrane | $\sim 7,500,000$ | 1 μs | 1.0 μs |
| Total length | | | | 529 μs |

### 2.1 Binding of closed-form mGBP2 to lipid membranes

To investigate how the closed form of mGBP2 interacts with lipid membranes, we performed unbiased 10 *µ*s CGMD simulations in triplicates with three starting orientations for the monomer across three membrane compositions: CL-mem (DOPC/CHOL/CL 75%/20%/5%), PS-mem (DOPC/CHOL/POPS 75%/20%/5%), and PC-mem (DOPC/CHOL 80%/20%).

PS-mem and PC-mem serve as control membranes, consistent with prior lipid-strip and GUV experiments showing reduced mGBP2 interactions with these compositions,^7,43,44^ while CL-mem represents the biologically relevant target membrane enriched in cardiolipin. The closed-form dimer was additionally simulated on CL-mem. In total, this yielded 27 monomer and 3 dimer simulations (**Fig. 1A,B**, **Table 1**), which we use to characterize the binding position and lipid selectivity of mGBP2, the influence of dimerization and starting orientation on membrane association, and the effect of membrane-bound mGBP2 on local lipid dynamics.

#### The mGBP2 features different membrane-binding modes, but binding via its geranylgeranyl anchor is the most stable

Across 27 monomer simulations, mGBP2 ends up inserted into the membrane via its geranylgeranyl anchor in 18 cases (binding pose 1, BP1; **Fig. 2A,B**). Two alternative binding poses were also observed: via the MiD/ED transition region (BP2, 6 cases) and via the GTPase domain (BP3, 1 case). In the remaining 2 simulations, mGBP2 did not associate with the membrane. Three simulations showed transient dissociation followed by rebinding.

**Figure 2:**
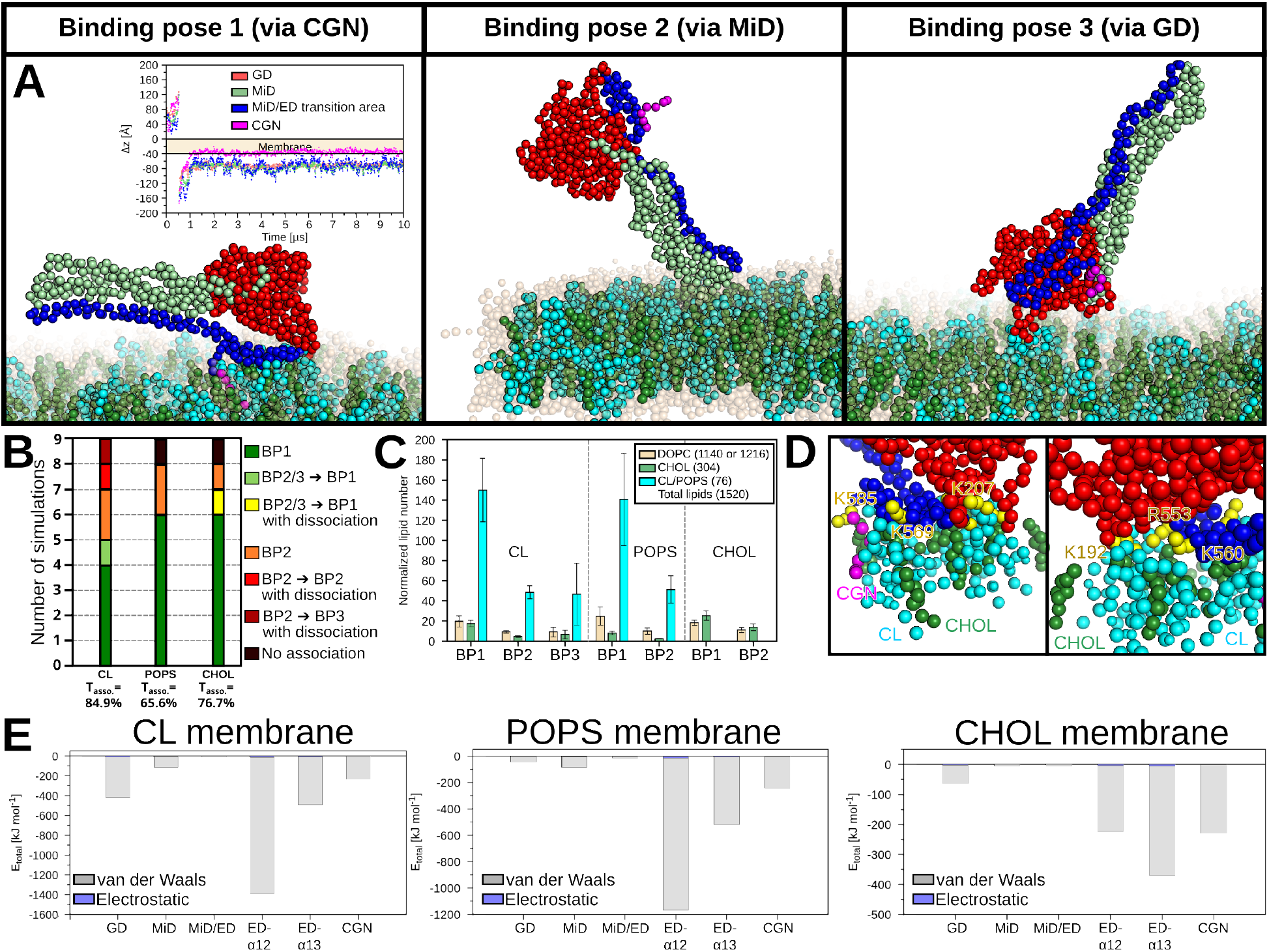
Results of the monomer systems of closed mGBP2. **(A)** During the unbiased CGMD simulations, three binding poses (BP) were identified: via (1) the geranylgeranyl group, (2) the MiD–ED transition region, and (3) the GD. Here, we show the most populated cluster for each binding pose. For the expected binding pose, we also depicted the time evolution of the Δ*z* position, with the membrane shown. **(B)** Statistical analysis of mGBP2 on the three different membranes (75% DOPC/20% CHOL with 5% CL or POPS as well as 80% DOPC/ 20% CHOL), nine simulations per membrane, 27 in total. Here, binding in BP1 is shown in green; binding in BP2/3 to BP1 without dissociation in bright green and with dissociation in yellow; binding in BP2 in orange; binding in BP2 with dissociation again in BP2 in red and in BP3 in dark red; and no association in black. Furthermore, the mean time of association (*T_asso._*) is computed across all nine simulations for each membrane. **(C)** For each membrane composition, the weighted average of the normalized lipid number was calculated for each binding poses (BP1-BP3). The weight was set to the number of frames where mGBP2 was binding in the specific binding pose. The normalized lipid number was determined by division of the lipid number and (lipid type number)/(total number of lipids). **(D)** A view of the interaction with CL is shown. **(E)** Energy profiles for the VDW (in gray) and Coulomb interactions (in blue) for each domain across the three tested membranes for all lipids. The examples were chosen based on the longest association time, and the single energy profile for each lipid is shown in **Fig. S4**.

Breaking this down by membrane composition, mGBP2 binds CL-mem predominantly in BP1 (5 of 9 cases) but also adopts BP2 (3 cases) or BP3 (1 case), and two of these runs involve dissociation and rebinding. On PS-mem and PC-mem, BP1 is even more dominant (6 and 7 of 9 cases, respectively), with one simulation showing no association on each membrane and one dissociation event on PC-mem, after which mGBP2 rebinds in BP1.

The membrane-association probability — defined as the average percentage of simulation frames in which the minimum protein–membrane distance remains below 0.5 nm — follows the order CL-mem (84.5%) *>* PC-mem (76.7%) *>* PS-mem (65.6%). Membrane association as such is therefore not selective in our simulations: mGBP2 also binds the two control membranes, and BP1 is even more frequent there than on CL-mem. This is not unexpected, as the hydrophobic geranylgeranyl anchor can insert into any bilayer once the protein is close to the membrane surface, and the simulations start with mGBP2 only 4 nm above the bilayer. The preference for CL becomes apparent instead in the nature of the contacts, as shown below: CL forms specific residue contacts and clusters at the binding site, whereas POPS contacts remain diffuse. Repulsion between polar or charged surface areas of mGBP2 and the membrane (**Figs. S3, S4**) accounts for the dissociation events observed on CL-mem and PC-mem.

BP1 is also the most stable pose in terms of membrane contacts: the normalized lipid contact number in BP1 exceeds 140 for both CL and POPS, whereas in BP2 this drops to approximately 50 — a roughly three-fold reduction (**Fig. 2C**). By contrast, DOPC and CHOL contacts remain low across all membranes (10–25 and 2.5–25, respectively), confirming that CL and POPS drive membrane association rather than the bulk phosphatidylcholine or cholesterol components. Once mGBP2 associates in BP1, it does not dissociate, as the geranylgeranyl anchor remains stably inserted in the membrane (**Fig. 2A**).

#### Residue-level interactions between mGBP2 and membranes

Membrane interactions are dominated by lysine and arginine residues, with the geranylgeranyl anchor consistently showing the strongest association (**Table S2**). Among polar residues, K560, K568, and K585 in the ED and K192 and R231 in the GTPase domain contribute most frequently. Notably, these ED lysines are not equivalent to the triple arginine motif (R584–R586) in hGBP1, which is positioned immediately adjacent to the farnesyl tail and forms a compact basic cluster. Depending on the extent to which the ED-*α*12 helix inserts into the membrane, the normalized lipid contact number increases by approximately 33% (**Fig. S4A**).

CL and POPS interact with mGBP2 in qualitatively different ways despite similar total lipid contact numbers. CL forms specific contacts with 4–46 residues, whereas POPS contacts are diffuse and largely non-specific (0–7 residues; **Table S2, Fig. S4B**). As expected from its abundance, DOPC accounts for the largest number of contacts (6.4–29.6 per lipid on average), while CHOL contributes fewer (0.4–6.4). CL and CHOL interactions are concentrated at lysine and arginine residues, as shown in **Fig. 2D**. Lipid density maps further support the specificity of CL interactions: CL and CHOL cluster at the protein binding site, whereas POPS shows no comparable enrichment (**Fig. S5**). CL clustering is also observed in protein-free reference simulations, though less pronounced than in the presence of mGBP2 (**Fig. S6**). The side-view density maps additionally show more pronounced membrane undulations near the mGBP2 binding site than in the protein-free references, most visibly on CL-mem, although we did not quantify this effect further.

We illustrate how the domain contributions vary using one representative trajectory per membrane, selected by the longest association time (**Figs. 2E** **and S4**). On CL-mem, the GD, geranylgeranyl anchor, and ED-*α*12 all contribute to binding. At the GD, DOPC and CL contribute similar interaction energies (DOPC: *−*211.4 kJ/mol, CL: *−*186.2 kJ/mol), although DOPC makes up 75% of the membrane and CL only 5%, while CHOL engages the ED and geranylgeranyl anchor with a strength comparable to CL. On PS-mem, the GD is largely uninvolved, but ED-*α*12 engages along its entire length with MiD also contributing, which brings the total non-bonded energy to *−*1166 kJ/mol, about three times more favorable than the *∼ −*370 kJ/mol of the other two examples. This extensive ED-*α*12 engagement is not specific to PS-mem but represents a binding mode that can occur on any membrane composition. On PC-mem, the interaction energies of DOPC and CHOL at ED-*α*12 and the geranylgeranyl anchor are comparable to those on CL-mem, and the total binding energy is similar despite the absence of CL.

As the total energy depends strongly on the binding mode and these are single representative trajectories, it does not allow us to rank the membranes. Instead, the role of CL is reflected in its high interaction energy relative to its abundance and in the specific residue contacts described above. POPS, in contrast, interacts much more weakly, although it is present at the same fraction of 5%: at ED-*α*12, its interaction energy is about *−*50 kJ/-mol, compared with *−*216.0 kJ/mol for CL, i.e., about four times weaker (**Fig. S4C**), even though the normalized contact numbers of the two lipids are similar. Together with the diffuse, largely non-specific POPS contacts, this is consistent with GUV experiments showing no significant mGBP2 accumulation on POPS-containing membranes.^32^

### The closed-state mGBP2 dimer binds membranes via a single geranylgeranyl anchor without adopting open-state conformations

The mGBP2 dimer can insert into the membrane via the geranylgeranyl anchor, but only through one protomer, as the anchor of the second protomer points away from the membrane in the closed-state dimer geometry (**Fig. S7A,B**). The dimer adopts the same binding poses and key-residue contacts as the monomer (**Fig. S7C**), but reaches a stable membrane-bound state more slowly: in all three replicates, which differed in the starting orientation of the dimer, this took 4.4–7.0 *µ*s, compared with typically less than 1–2 *µ*s for the monomer (**Table S1**). Simultaneous two-anchor membrane binding would require large conformational changes, which are not observed in our CGMD simulations of the closed, nucleotide-free state. For hGBP1, such opening requires nucleotide turnover rather than nucleotide binding alone: while GTP binding suffices for binding to host-like vesicles, polymer formation requires GTP hydrolysis;^45^ full-length hGBP1 dimerizes only in the presence of the hydrolysis transition-state mimic GDP *·* AlF_3_,^23^ and coat formation on bacteria requires continuous GTP hydrolysis, as transition-state mimics and non-hydrolysable analogues fail to produce coats. ^30^

### 2.2 The open-state GBPs in solution

The membrane-bound conformations of the closed mGBP2 dimer, with only one prenyl anchor inserted and the dimer contacting the membrane via the MiD/ED transition region (Section 2.1), cannot be reconciled with the arrangement of upright, doubly anchored dimers in hGBP1 coats resolved by cryo-electron tomography.^23,30^ In these coats, the individual proteins are extended to a length of up to 28–29 nm, whereas closed GBPs measure only about 13 nm. Coat formation thus requires the open state, which has been characterized for hGBP1 but not for mGBP2. Given the high sequence identity between the two proteins, we modeled open monomeric and dimeric mGBP2 structures and assessed their stability in solution by CGMD simulations of the prenylated (holo) proteins. For the dimer, we did not use the AlphaFold 3 prediction (**Fig. S1**), as it lacks the criss-cross arrangement at the GD–MiD transition region; without it, the two protomers separated at the level of their GDs and MiDs within less than 5 *µ*s in our previous simulations.^28^ Instead, we built the mGBP2 dimer by homology modeling with SwissModel,^34^ based on the criss-cross hGBP1 dimer (PDB: 8CQB).^23^ Although no nucleotide was included, the nucleotide-binding region of the GD was restrained to the conformation of the template structures, which were solved with the hydrolysis transition-state mimic GDP *·* AlF_3_,^23,25^ so that the simulations represent the nucleotide-activated state. In previous work, we applied the same approach to open monomers and dimers of hGBP1 and hGBP5.^28^ Here, we extend it to an open hGBP1 hexamer, an assembly of three dimers resolved by cryo-ET of hGBP1 (PDB: 8R1A),^22^ which represents the next step towards polymeric coats.

#### Dimerization stabilizes the open state of mGBP2

We assessed the stability of the open structures by comparing start and end structures and by the ED–GD distance (**Fig. 3A,B**), calculated between residue 582 at the tip of the ED and residue 312 in the GD. The open mGBP2 monomer shows the highest flexibility: the ED–GD distance reveals that the extended ED can transiently fold in toward the GD before reopening, which occurs repeatedly during the simulation. However, the spatial distribution of residue 582 shows that the closing is not complete, i.e., the ED tip does not reach the GD (**Fig. S8A**). The root mean square deviation (RMSD) (**Fig. S8B**) further confirms that the GD and MiD remain stable throughout and only the ED undergoes these large-scale motions.

**Figure 3:**
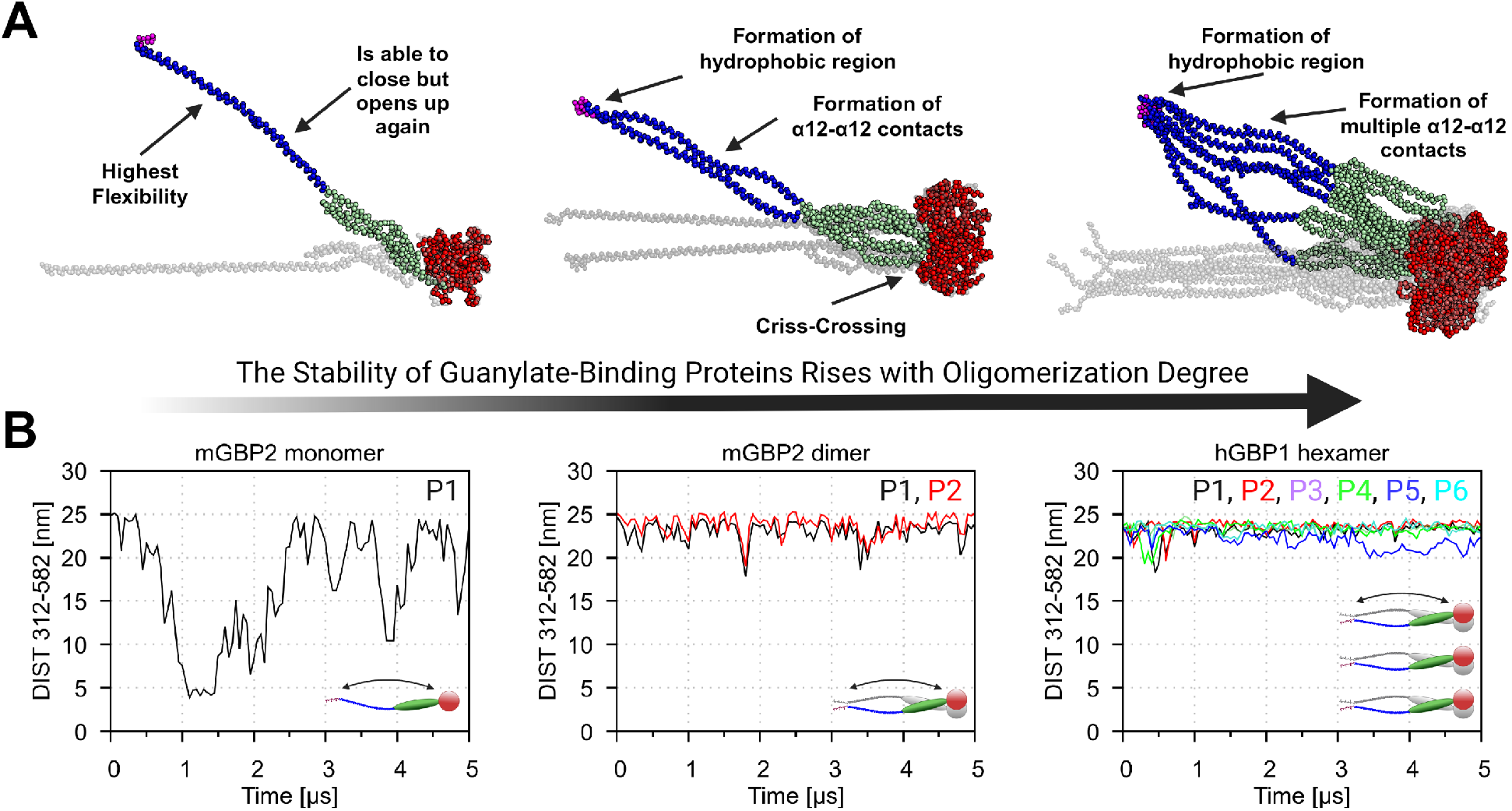
Results of the open GBPs in solution. **(A)** The structures before (gray) and after 5 *µ*s CGMD simulations of the open mGBP2 monomer, mGBP2 dimer, and hGBP1 hexamer in solution are presented (GD in red, MiD in green, ED in blue, and prenylation in magenta). Striking differences between the monomer, the dimer and the hexamer are annotated. **(B)** The minimal distances between residues 312 to 582 recorded during 5 *µ*s CGMD simulations of the open mGBP2 monomer, mGBP2 dimer, and hGBP1 hexamer in solution. The individual protomers are labeled (P1 to P6) and an in-plot cartoon illustrates the measured distance for tracking possible ED-to-GD closing in the GBPs.

The open mGBP2 dimer is considerably more stable than the monomer. The criss-cross interface at the GD-to-MiD transition point enables additional *α*12–*α*12 interactions — which could lead to coiled-coil formation, as characterized in detail for hGBP1 and hGBP5^28^ — and limits motions of the EDs compared to the monomer. No transient closing is observed (**Fig. 3B**), and the averaged RMSD of the dimer remains below 4 nm (mostly below 2 nm), compared with values of up to 8 nm for the monomer (**Fig. S8B**). In particular, the motion at the MiD-to-ED transition is reduced, as the spatial distribution of residue 480 shows (**Fig. S8A**).

#### A hexamer further stabilizes open hGBP1 dimers

The open hGBP1 hexamer shows the lowest ED–GD distance fluctuations among the three systems and lower than those of isolated hGBP1 dimers in our previous work^28^ (**Fig. 3B**). The central dimer is the most stable, as it is flanked by two additional dimers that further restrict its movement. Additional stabilization of the hexamer arises from *α*12–*α*12 interactions and from the clustering of the six farnesyl anchors into a hydrophobic region at one end of the assembly, which, together with the larger size of the GDs at the other end, gives the hexamer a cone-like shape.

Taken together, dimerization stabilizes the open state of mGBP2 in the same way as previously shown for hGBP1 and hGBP5,^28^ and assembly of three dimers into a hexamer stabilizes the open hGBP1 dimers further. Building on these solution-phase findings, we next examined whether the same stabilization principles hold when open GBPs encounter lipid membranes.

### 2.3 The open-state GBPs at membranes

In this part of the study we investigate the stability of open hGBP1 and mGBP2 structures at different membrane interfaces and their effects on the bilayer. Starting structures are shown in **Fig. 1C,D,E**; all simulations are summarized in **Table 1**. We address two central questions: how does protein–membrane interaction change when switching from closed to open states with pre-inserted prenyl groups, and how do hGBP1 and mGBP2 stabilize their open conformations at membranes?

To study the open conformation of hGBPs and mGBPs at the membrane interface, we selected asymmetric bilayer compositions to mimic biological membranes; the lipids used are shown in **Fig. S2**. Three membrane systems were used. First, a pure POPC bilayer served as a simple reference for both hGBP1 and mGBP2 simulations. Second, a biologically realistic LPS membrane was constructed, with 80% LPS and 20% POPE in the outer leaflet, and 90% POPE, 5% 1-palmitoyl-2-oleoyl-*sn*-glycero-3-phosphoglycerol (POPG), and 5% cardiolipin (CL) in the inner leaflet, matching the asymmetric outer membrane composition of Gramnegative bacteria;^46–48^ this system was used exclusively for hGBP1 simulations. The LPS fraction of 80% was chosen based on protein-free titration simulations with a simplified POPC-based bilayer and 30–100% LPS in the outer leaflet (**Fig. S9**), in which 80% LPS already yielded a well-covered outer leaflet, and on the fact that LPS does not cover the entire outer leaflet of Gram-negative bacteria.^49,50^ Third, a PI5P membrane motivated by recent findings on the lipid composition of pathogen-containing vacuoles and mGBP2 lipid specificity^32,51–54^ was constructed with 5% PI5P, 20% cholesterol (CHOL), and 75% DOPC in the outer leaflet, and 20% CHOL and 80% DOPC in the inner leaflet; this system was used exclusively for open mGBP2 simulations. In the following, these systems are referred to as the POPC, LPS, and PI5P membranes, respectively.

#### Lipid environment shapes the conformational dynamics of open-state GBPs

Since open GBPs are anchored to the membrane via their prenyl group, the key question is how the open conformation behaves once membrane-bound and whether the lipid environment can stabilize it. We therefore compared representative snapshots of the binding modes and GD–membrane distances for monomers and dimers of both species (**Fig. 4A** and **Fig. S10**). The dimer is included because it is the smallest assembly unit that forms the basis for polymeric coat formation.

**Figure 4:**
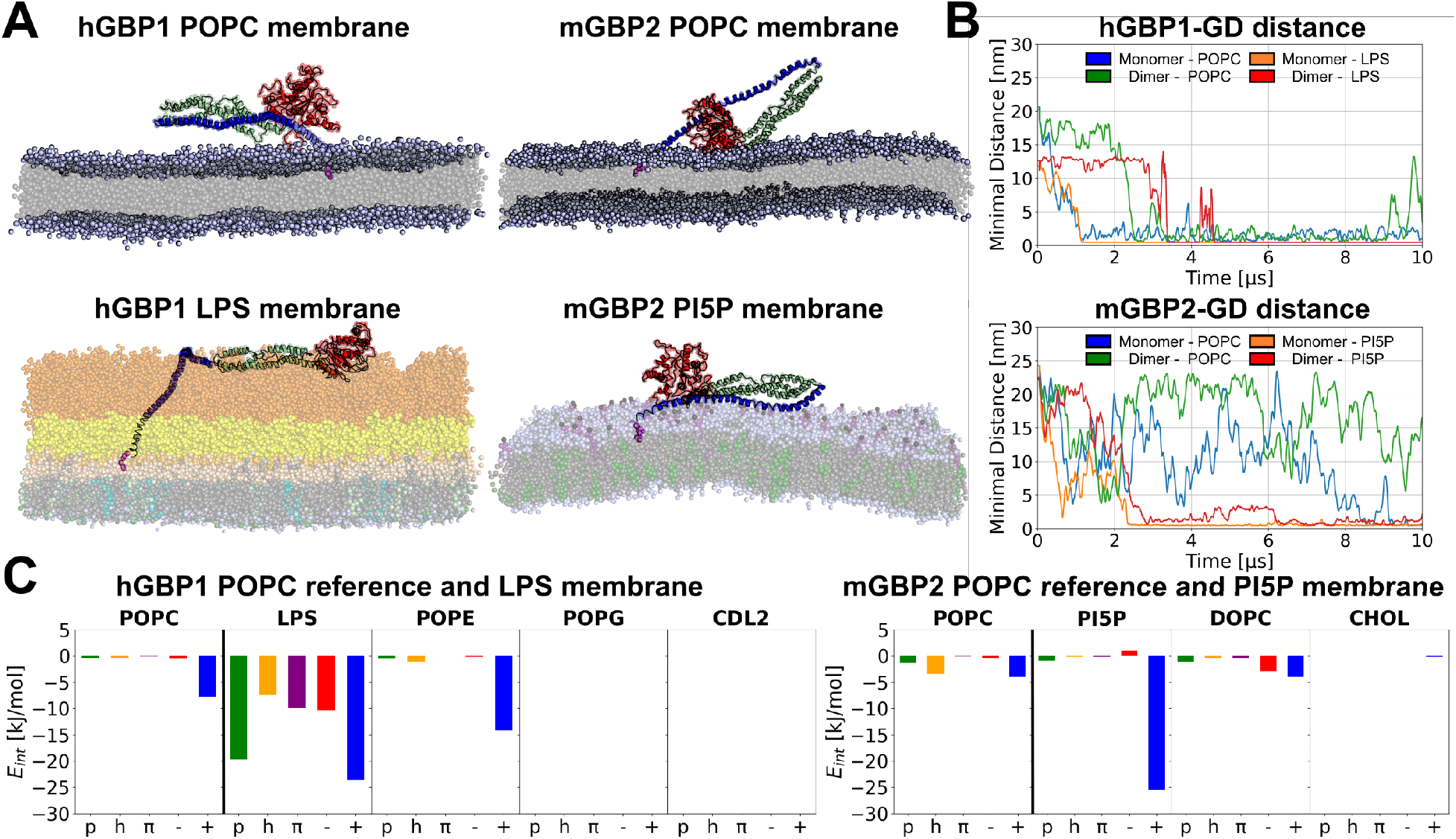
Results of the open GBPs on membranes. **(A)** Backmapped representative snapshots from the CGMD simulations of monomeric hGBP1 and mGBP2 on membranes. The POPC membranes (top) serve as reference. The LPS membrane (bottom left) is composed of 80% LPS and 20% POPE in the upper leaflet and 90% POPE, 5% POPG, and 5% CL in the lower leaflet. The PI5P membrane (bottom right) is composed of 75% DOPC, 20% CHOL, and 5% PI5P in the upper leaflet and 80% DOPC and 20% CHOL in the lower leaflet. Snapshots of the dimeric hGBP1 and mGBP2 membrane systems can be found in the Supporting Information (**Fig. S10**). **(B)** Minimal distance between the GDs and the membrane component of interest: POPC for the reference simulations with hGBP1 and mGBP2, LPS for hGBP1, and PI5P for mGBP2 monomers and dimers. The distance are shown for GBP monomers and dimers as running averages. The different starting distances are due to the different outer leaflet thickness. **(C)** Energetic analysis of monomeric GBP interaction with each membrane component. The interactions with POPC result from the reference simulations. The interaction energies (*E_int_*) were normalized with respect to the number of residues contacting each lipid and are shown for the protein residues grouped by amino acid type (polar: p; apolar: h; aromatic: *π*; negatively charged: *−*; positively charged: +).

In all simulations, the GD–membrane distance decreases (**Fig. 4B**), and all GBP monomers and dimers increase their membrane contact area, though the mechanism differs between species and membrane type. On the POPC membrane, monomeric hGBP1 folds back fully within the simulation time and reaches a closed state that resembles the available crystal structures,^55,56^ but with the ED on the opposite side of the GD and MiD (see below). In the hGBP1 dimer, no hinge motion at the MiD–ED transition takes place; instead, the EDs bend, bringing the MiDs and GDs toward the membrane surface (**Fig. S10**). On the LPS membrane, hGBP1 also does not fold fully, and the O-antigen chains shape the binding mode of the monomer and the dimer differently. The monomer inserts the lower part of its ED into the O-antigen layer and kinks at the position where the O-antigen chains end. The remaining part of the protein, comprising the rest of the ED, the MiD, and the GD, lies flat on top of the O-antigen layer. In the dimer, the EDs bend without forming a kink, so that only the GDs come into contact with the LPS.

For mGBP2, hinge-driven folding at the MiD–ED transition is observed for the monomer on both the POPC and PI5P membranes, but the outcomes differ markedly. On the POPC membrane, the hinge motion leads to membrane adsorption on the GD/MiD side with the ED pointing away from the membrane, yet without reaching stable attachment within the simulation time. On the PI5P membrane, by contrast, mGBP2 closes fully and substantially increases its membrane contact area via the ED, which is consistent with a strong electrostatic attraction between positively charged surface patches of mGBP2 and the negatively charged inositol phosphate head groups of PI5P, as detailed below. Similar to the hGBP1 dimers, mGBP2 dimers do not undergo closing via MiD–ED hinging; instead, both dimers remain in a largely extended conformation (**Fig. S10**) and tilt toward the membrane surface through conformational changes in the C-terminal region near the geranylgeranyl anchor. On the POPC membrane, the mGBP2 dimer hovers above the membrane without stable contact, whereas on the PI5P membrane it lies flat against the surface, again pointing to the importance of electrostatic interactions with PI5P head groups.

Overall, the dimers of both species remain largely extended, whereas the monomers fold back or kink, preferentially via the MiD–ED region. This agrees with the higher stability of the open state in dimers than in monomers observed in solution, which is a consequence of criss-crossing at the GD–MiD transition region leading to stabilizing interprotein interactions.^28^ The EDs of the dimers nonetheless adapt to the membrane. Their end-to-end distance, measured between residues 480 and 582, decreases from 15.3 nm in the starting structure to 12.5 *±* 0.3 nm for hGBP1 on POPC and 10.4 *±* 0.3 nm on the LPS membrane, whereas for mGBP2 it remains close to the starting value of 16.4 nm, at 15.3 *±* 0.1 nm on POPC and 15.1 *±* 0.1 nm on the PI5P membrane. The EDs of hGBP1 are thus considerably more deformable than those of mGBP2, and the two membranes act on them differently: on the LPS membrane the GDs are drawn towards the O-antigen layer, bending the EDs, whereas on POPC the shortening reflects the intrinsic flexibility of the EDs in the absence of such attraction.

#### Binding-site analysis localizes lipid contacts on the protein

To determine where on the protein the membrane contacts occur, we performed PyLipID binding-site analysis for all monomers and dimers (**Fig. S11**) and labeled the top three protein residues for membrane interactions per system. On the POPC membrane, contacts are sparse and differ between species and oligomeric states. The hGBP1 monomer contacts the bilayer mainly along *α*13 of the ED, with some additional contacts of the GD, whereas the hGBP1 dimer contacts it mainly via the C-terminus, with only a few, less frequent contacts of the GD. The mGBP2 monomer contacts POPC mainly through the GD and the GD–MiD transition region, in addition to the C-terminal end, and the mGBP2 dimer shows only C-terminal contacts and short-lived contacts of the GD. On the LPS membrane, hGBP1 contacts extend along the ED up to the reach of the O-antigen chains. The monomer additionally contacts the MiD– ED transition region, and both the monomer and the dimer contact the GD. On the PI5P membrane, both the mGBP2 monomer and dimer form binding sites along the entire protein. The dimer shows a larger number of binding sites along the MiDs and EDs, consistent with it lying flat on the membrane in the open conformation. Thus, LPS and PI5P extend the contact surface well beyond the lipid anchor, whereas the contacts on POPC are sparse and, for the dimers, largely restricted to the C-terminus. Density maps show that PI5P is enriched around regions of high protein density (**Fig. S11D,H**), even though it makes up only 5% of the upper leaflet. LPS, which makes up 80% of the upper leaflet, shows a less pronounced reordering, but also accumulates around hGBP1.

Per-residue association times (**Figs. S12 and S13**), which include only residues in contact for at least 10% of the simulation time, identify the most persistent contacts. These are dominated by the lipid anchor and its neighboring residues, which engage the membrane from the start, whereas contacts further along the protein form only later in the simulations and therefore remain largely below this threshold. On the LPS membrane, residues along the entire hGBP1 stalk nevertheless engage LPS persistently, whereas POPC and POPE mainly contact the C-terminal polybasic motif and adjacent lysines, and POPG and CL contact the farnesyl anchor only briefly. For mGBP2, persistent contacts with POPC are likewise restricted to the C-terminal residues and the geranylgeranyl anchor, while on the PI5P membrane, residues of the GD and ED also interact persistently with PI5P and DOPC.

#### Positively charged residues drive GBP–membrane interactions

To understand these driving forces in detail, we decomposed the protein–membrane interaction energies of the GBP monomers by residue type into contributions from positively charged, negatively charged, polar, apolar, and aromatic residues (**Fig. 4C**). Because the open hGBP1 and mGBP2 monomers were simulated with MARTINI 2.2 and MARTINI 3, respectively, we compare absolute energies only within, not across, species. For hGBP1, interactions with the POPC membrane are weak and mediated by only a small number of positively charged residues, whereas in the LPS membrane, interactions with both POPE and LPS are substantially stronger and likewise dominated by positively charged residues. For LPS in particular, polar, apolar, aromatic, and negatively charged residues also contribute notably, reflecting the multitude of chemical moieties presented by its oligosaccharide chains. The overall dominance of positively charged residues across POPE and LPS is consistent with a strong electrostatic attraction of hGBP1 to the negatively charged outer membrane of Gram-negative pathogens. POPG and CL, located in the inner leaflet, show no interactions with protein residues; CL contacts only the inserted farnesyl anchor (**Fig. S14**). For mGBP2, a similar pattern emerges: interactions with the POPC membrane are weak, while the presence of PI5P strongly increases the interaction strength, driven predominantly by positively charged residues attracted to the negatively charged inositol phosphate head groups. DOPC, as the bulk structural lipid in the PI5P membrane, shows a more balanced interaction profile, with both positively and negatively charged residues contributing on a per-residue basis. Cholesterol exhibits the fewest interactions with mGBP2, reflecting its deeper position in the membrane core and the shielding of its small head group by the larger head groups of PI5P and DOPC.

The per-lipid and per-residue breakdown, which also includes the prenyl anchors that are not part of the residue-type decomposition in **Fig. 4C**, shows the same pattern for both species (**Figs. S14, S15, and S16**): the characteristic lipid of each membrane accounts for most of the interaction, whereas the other lipids interact mainly with the C-terminal anchor region, i.e., the prenyl anchor and adjacent basic residues. For hGBP1, the most favorable contributions to the LPS interaction arise from the C-terminal polybasic motif and adjacent lysines binding the phosphate groups of the LPS inner core and lipid A, complemented by many weaker, mainly Lennard-Jones contacts with the outer core and O-antigen; POPE interacts with the farnesyl anchor and the polybasic motif. For mGBP2, the PI5P interaction is mediated by lysines (K207, K212, K213, K243, K247, K581, K585) and arginines (R215, R242) of the GD and ED, whereas DOPC and CHOL interact mainly with the geranylgeranyl anchor and K585.

#### Open GBPs fold back into closed states that differ in ED orientation

Resting, inactive states of GBPs are predominantly described in the literature as ED-to-GD closed, nucleotide-free, or GDP-bound monomers, in which the geranylgeranyl and farnesyl lipid anchors are buried in hydrophobic pockets. ^16,32,57,58^ Human GBPs in resting states are largely cytosolic and soluble,^16^ while mGBPs can reside in vesicle-like structures in homo- and heterodimeric assemblies.^59^

In our current CGMD simulations of the open hGBP1 monomer on a pure POPC membrane, hGBP1 folds back at the MiD–ED transition region within 10 *µ*s and reaches a closed state. This state hovers above the membrane, with the inserted farnesyl anchor and the membrane-proximal residues of *α*13 as the main contacts, and with some additional contacts of the GD (**Fig. 4A**). A striking feature of this state is the switching of the ED from one side of the GD and MiD to the other (**Fig. S17A**), which is not observed in any available PDB structure. An alternative GD–ED docking site has, however, been identified by single-molecule FRET spectroscopy of hGBP1.^24^ Alternative ED binding sites are also accessible in other GBPs: in our previous MD simulations, holo-hGBP5 occupied an ED binding site involving helices *α*0/1 instead of the canonical site at helices *α*3/4.^28^ The new ED position of hGBP1 is supported by intramolecular contact matrices (**Fig. S17A**). A second striking feature is the behavior of *α*13: whereas in all known PDB structures *α*13 folds back against *α*12 as a separate helix, here *α*13 forms a continuous extension of *α*12, creating one elongated helix. This is consistent with our earlier CGMD work, which identified *α*13 as highly flexible and prone to detaching from *α*12,^58^ and it is this elongated configuration that allows hGBP1 to hover above the membrane. It also explains why the folded-back ED resembles the crystal structures only in a similar, but not identical, position.

This arrangement corresponds to the conformer M2 identified for hGBP1 in solution by integrative structural biology, in which *α*12/13 binds the side of the GD opposite to that seen in the crystal structures.^24^ Superimposing our closed hGBP1 structure on the 206 conformers of the deposited M1/M2 ensemble (PDBDEV 00000088) gives a minimum backbone RMSD of 4.8 Å to an M2-type conformer. These conformers were generated using FRET and DEER distances as restraints and carry a corresponding structural uncertainty, with root-mean-square fluctuations of about 7–9 Å within each ensemble and the lowest precision in the *α*12/13 region;^24^ our structure therefore agrees with the M2 conformer to within the resolution of that ensemble. Our simulations thus show that hGBP1 can adopt this conformation while remaining membrane-bound through its inserted farnesyl anchor.

For mGBP2, a similar fold-back via the MiD–ED hinge occurs within 10 *µ*s on asymmetric PI5P membranes, with *α*12 and *α*13 likewise forming one elongated helix and the geranyl-geranyl anchor deeply embedded. In contrast to hGBP1, this closed state is membrane-engaged rather than hovering, and the ED does not switch sides but retains an orientation consistent with the AlphaFold model of mGBP2, which, given 68% sequence identity with hGBP1, is nearly identical in fold to hGBP1. This is further supported by contact maps and by comparison with our previous atomistic simulations of mGBP2 membrane interaction ^32^ (**Fig. S17B**). Notably, the ED engages directly with PI5P head groups, which may stabilize this orientation. Thus, both proteins fold back into closed states with an elongated *α*12/13 helix, but with opposite ED orientations, which, however, formed on different membranes (POPC for hGBP1, PI5P for mGBP2).

#### Long O-antigen chains keep open hGBP1 upright

The LPS membranes described above carry the longest O-antigen chains available in the CG model, which extend about 5 nm above the membrane and thus reach only to about the middle of the ED. We therefore asked whether longer chains, with 12 repeat units matching the average chain length of *S. flexneri*,^61^ can keep open hGBP1 upright. Since no coarse-grained model for LPS with long O-antigen chains is available, we simulated this system with AAMD and restricted it to the hGBP1 dimer. The chains were built in an extended conformation reaching about 20 nm above the membrane and relaxed to about 14 nm during the simulation, comparable to the O-antigen extension of 10–13 nm measured for LPS from *S. enterica* and *E. coli* .^23^

On the long-chain membrane, the dimer remains upright for the full 2*×*1 *µ*s of simulation, and the GDs do not tip towards the membrane (**Fig. 5D-F**). As the O-antigen layer relaxes, the chains initially reach the GDs but by the end only the middle of the MiDs. Density maps confirm this (**Fig. 5E**) and additionally show flip-flop of POPG and CL towards the outer side of the membrane (blue circles) as well as a local convex deformation of the outer leaflet around the site of hGBP1 insertion, which we did not quantify further. PyLipID identifies LPS binding sites on both hGBP1 chains (**Fig. 5F**). LPS interacts more strongly with one of the two chains, as shown by larger spheres proportional to the residence times. The smaller spheres at the top of the MiD indicate less stable binding, in line with the sinking of the LPS layer. With residence times of up to 430 ns within the 1 *µ*s simulations, the contacts between hGBP1 and LPS are long-lived.

**Figure 5:**
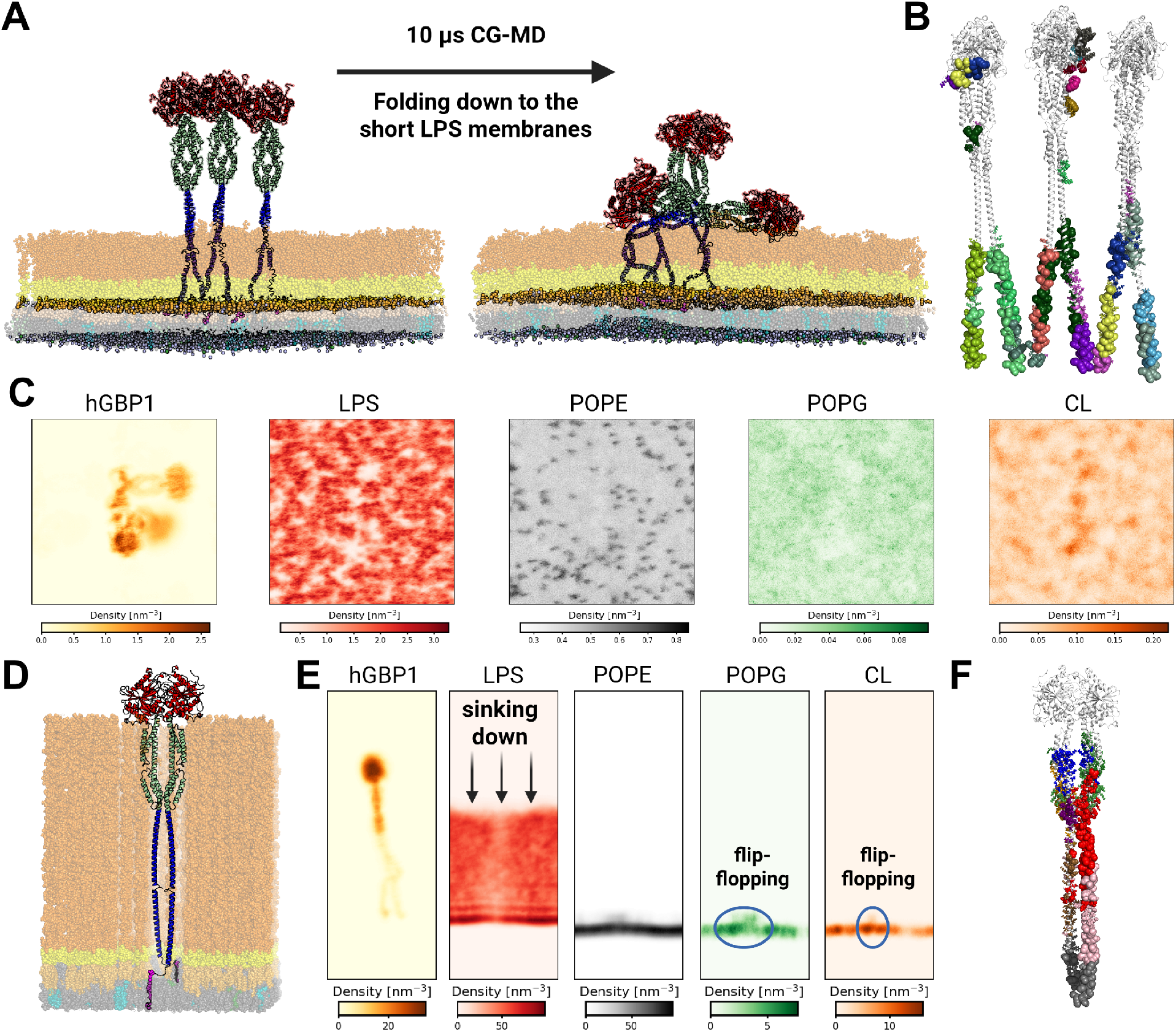
Open hGBP1 on LPS membranes with short and long O-antigen chains. **(A)** The hGBP1 hexamer on the short O-antigen chain LPS membrane before and after 10 *µ*s of CGMD simulation. The membrane is composed of 80% LPS and 20% POPE in the upper leaflet and 90% POPE, 5% POPG, and 5% CL in the lower leaflet. **(B)** Protein–LPS binding sites of the same run, calculated with PyLipID^60^ and projected onto the hGBP1 hexamer structure. **(C)** Top view of the density maps of all membrane and protein components. **(D)** Starting structure of the AAMD system with the hGBP1 dimer on the long O-antigen chain LPS membrane. The membrane composition is as in (A), but with 12 O-antigen repeat units and at all-atom resolution. **(E)** Density maps of all membrane and protein components. The relaxation of the O-antigen layer during the 1 *µ*s AAMD is marked, flip-flop events of POPG and CL are circled in blue, and the outer leaflet bulges outward around the site of hGBP1 insertion. **(F)** Protein–LPS binding sites of the hGBP1 dimer on the long O-antigen chain LPS membrane, calculated with PyLipID.

Short O-antigen chains, in contrast, do not hold the protein upright, and higher-order assembly does not compensate for this: the CG hGBP1 hexamer tips over towards the short-chain LPS membrane (**Fig. 5A-C**). Its three dimers face different directions, and only one remains upright (**Fig. 5A**). Because the long- and short-chain systems differ in oligomer size and resolution, we do not compare them quantitatively. Besides the prenyl anchor, which remains the main anchoring motif, hGBP1 forms several smaller LPS binding sites along the ED, in the MiD–ED transition region, and in the GD. In contrast, the POPC and PI5P membranes did not provide interfacial interactions strong enough to keep open GBP monomers and dimers in their initial upright placement.

### 2.4 GBP polymerization

To test whether higher degrees of oligomerization stabilize open hGBP1 on membranes, we simulated a hexamer, an octadecamer (18mer), and a periodic coat-like assembly on POPC membranes with MARTINI 3 (**Fig. 1F**, **Table 1**). We further simulated a disc of 30 hGBP1 dimers (60mer) to ask how a polymeric assembly engages a membrane. Because larger systems with realistic membranes were computationally out of reach, we used simple POPC membranes.

#### Higher-order assemblies reduce, but do not abolish, tipping towards the membrane

Membrane coats of GBPs consist of thousands of molecules. ^30^ To test how protein– protein interactions affect coat formation, we started with a hexamer, which consists of three hGBP1 dimers arranged in a row. In the 5 *µ*s simulation of the hGBP1 hexamer on a POPC membrane, one dimer left the hexameric and protein–membrane interface early in the trajectory, and its farnesyl anchors later re-inserted into the bilayer (**Fig. 6A**). The interface between the other two dimers remained intact, but the hexameric interface was not restored. However, all hGBP1 molecules, i.e. the single dimer and the remaining tetramer, remained in the open state. This contrasts with the hexamer on the LPS membrane described above. There, the hexamer stayed intact and did not leave the membrane, but two of its three dimers tipped over and did not return to the upright state (**Fig. 5A**). The O-antigen chains of LPS surround the helical stalks of the EDs. We interpret this as attractive interactions between the O-antigens and the protein regions above them (mainly the MiDs and GDs), which pull the stalks towards the membrane and kink the open hGBP1. Individual LPS molecules in contact with the protein are transiently displaced by about 0.2–0.3 nm, and local indentations of similar magnitude appear where the GD first contacts the O-antigen layer. It should be noted that both comparisons rest on single trajectories and on different force fields (MARTINI 2.2 for the LPS system, MARTINI 3 for the POPC system). As MARTINI 2 tends to overestimate protein–protein attraction, which was reduced in MAR-TINI 3,^39,62^ the greater cohesion of the hexamer on LPS may partly reflect the force field. We therefore regard these comparisons as tendencies.

**Figure 6:**
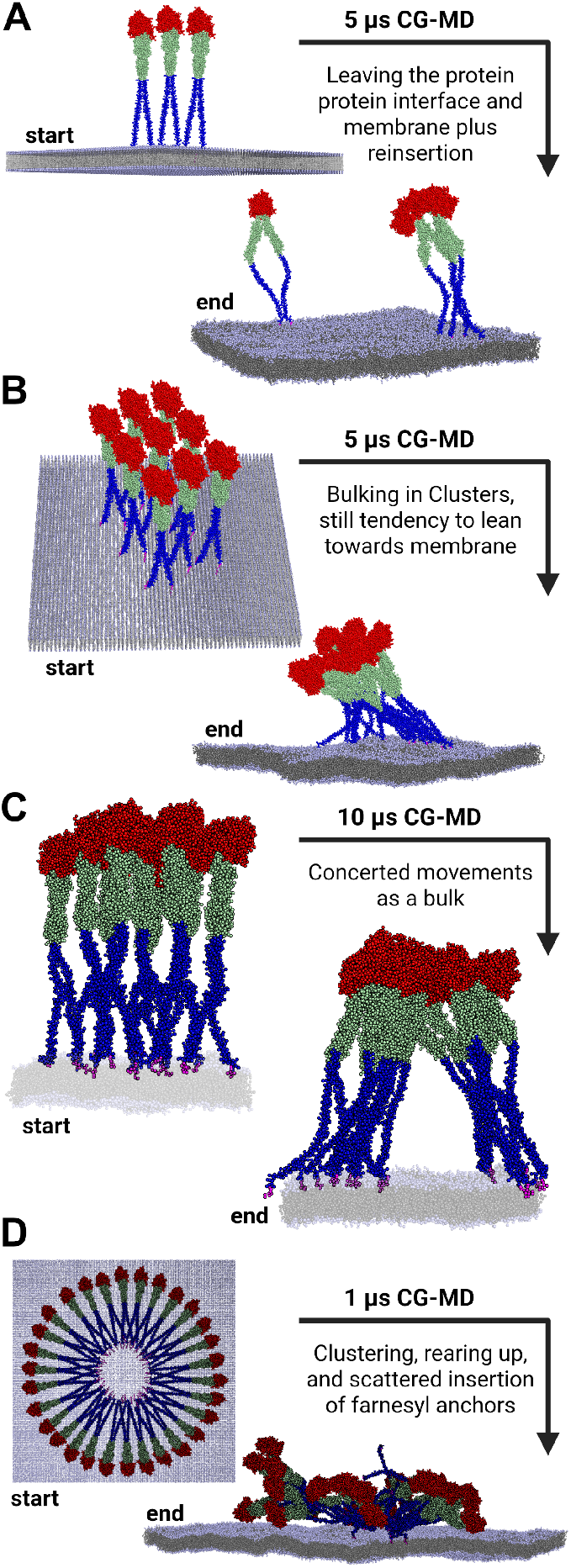
Higher-order hGBP1 assemblies on POPC. Each system is shown before and after the CGMD simulation, with the simulation time given and the main observations annotated. **(A)** The hGBP1 hexamer, from which one dimer dissociates. **(B)** The octadecamer, whose edge molecules arch towards the membrane. **(C)** The periodic coat-like assembly, in which all molecules remain upright. **(D)** The disc of 30 hGBP1 dimers during initial docking onto the membrane.

Next, we simulated an 18mer on a POPC membrane, which we constructed by placing three hexamers behind each other. The simulation time was again 5 *µ*s. The 18mer stayed intact, and the GDs clustered. Some of the farnesyl anchors moved away from each other within the membrane (**Fig. 6B**). None of the hGBP1 molecules kinked, i.e. none showed a hinge motion, but some of them, particularly those at the edges of the 18mer, arched towards the membrane. This motion was reversible within the simulation time, and no new resting state formed (**Fig. 6B**). These observations suggest that a larger assembly preserves the integrity of the oligomer on the membrane while the open state is retained. The 28–29 nm long hGBP1 molecules nevertheless remain flexible.

To mimic a coat, we placed an 18mer, i.e. three hexamers behind each other, on a POPC membrane such that it touched its periodic images in the membrane plane (*xy*). The resulting protein layer is thus effectively infinite in this plane (**Fig. 6C**). In contrast to the isolated 18mer on the larger POPC membrane, all hGBP1 molecules remained upright and open, as they were confined laterally by their neighbors. The bulky GDs and MiDs of neighboring molecules pack against each other, with the GDs interacting via their outer helices and loops, whereas the thin, single-helix EDs leave a void between them near the membrane. On the POPC membrane, this void is filled with water, and two subclusters of hGBP1 leaned slightly against each other. The void thus lies in the ED region, i.e., at the height that the short O-antigen chains of our LPS model reach (Section 2.3). Taken together, lateral confinement within an extended coat, which the isolated hexamer and 18mer lack, suffices to keep open hGBP1 upright, and the only remaining flexibility arises from the void between the EDs.

#### An hGBP1 disc starts to engage a POPC membrane

To model a single layer of the polymeric storage form, ^22,23^ we placed a disc of 30 hGBP1 dimers about 4 nm above a POPC membrane (100 *×* 100 nm^2^, approx. 7.5 million beads) and simulated it for 1 *µ*s. The disc separated into smaller oligomeric clusters of up to five dimers. Chains of clusters tilted away from the membrane, and a few farnesyl anchors inserted into the bilayer (**Fig. 6D**). The farnesyl anchors are spaced widely in the disc. We speculate that this spacing prevents joint movement of the whole disc and that denser packing might favor it. Because this is a single 1 *µ*s trajectory, we regard it as a snapshot of the initial phase of membrane docking. Across the larger assemblies, side-view density maps show membrane undulations beneath the bound proteins (**Fig. S18**).

## 3 Discussion

How GBPs engage pathogen membranes has so far been studied either at the level of isolated proteins in solution or at the level of entire coats on bacteria, leaving the intermediate steps largely inaccessible. Molecular dynamics simulations have addressed the closed, resting states of GBPs and related dynamin-like proteins,^32,58,63,64^ but the open, membrane-active states have not been examined at membranes. Here we simulated closed and open hGBP1 and mGBP2 as monomers, dimers, and higher-order assemblies up to up to a disc of 30 hGBP1 dimers, on membranes ranging from pure phosphatidylcholine to *S. flexneri* LPS and phosphoinositide-containing bilayers, covering more than 0.5 ms (over 500 *µ*s) of cumulative sampling. Three findings emerge, which together with the steps leading up to them are summarized in **Fig. 7**: the open state is intrinsically unstable at the monomer level and is maintained by oligomerization; open GBPs fold back into distinct membrane-bound resting states that differ between the two species; and keeping an open GBP upright on a membrane requires either long O-antigen chains or the lateral confinement of an extended coat.

**Figure 7:**
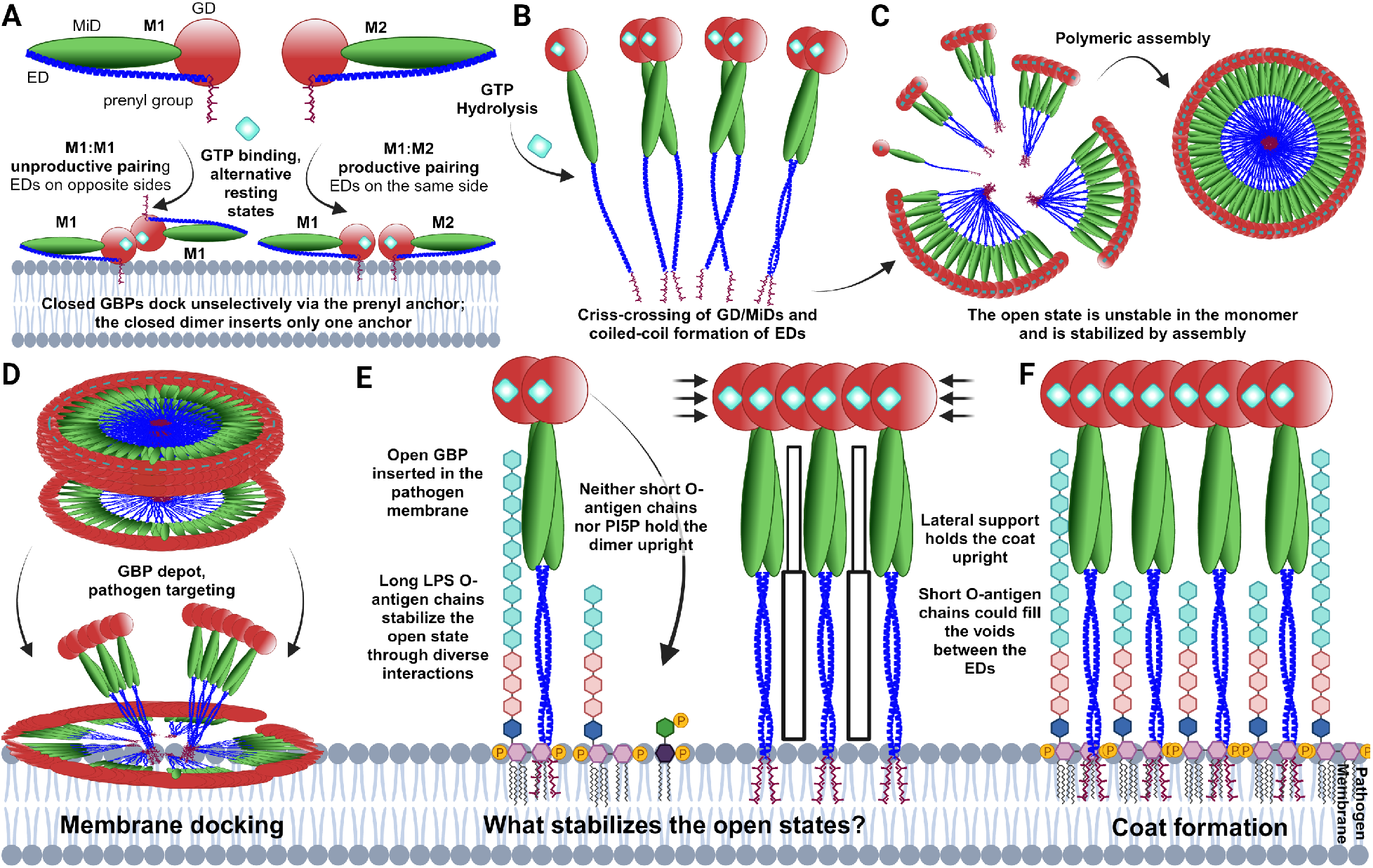
Model of GBP membrane engagement emerging from our simulations. **(A)** Closed GBPs dock to membranes through their prenyl anchor, largely irrespective of lipid composition, and the closed dimer inserts only one of its two anchors. Open GBPs that fold back at a membrane adopt alternative resting states with the anchor remaining inserted, corresponding to the conformers M1 and M2;^24^ only an M1:M2 pairing places both EDs on the same side and can open again. **(B)** Nucleotide binding and hydrolysis open the protein and drive dimerization through criss-crossing of the GD–MiD regions and coiled-coil formation of the EDs. **(C)** In solution, the open monomer is unstable and folds back, whereas the open state is stabilized by assembly; in the absence of lipids, GBPs form a polymeric assembly. **(D)** This polymeric form acts as a depot: near the membrane it dissociates into smaller clusters from which individual prenyl anchors insert. **(E)** What keeps an open GBP upright at a membrane: long O-antigen chains reaching into the MiD region stabilize the open state, whereas neither short O-antigen chains nor PI5P hold the dimer upright; within a laterally confined coat, neighboring molecules support each other. **(F)** In the coat, the bulky GDs and MiDs of neighboring molecules pack against each other, leaving voids between the thin EDs close to the membrane that short O-antigen chains could fill. GBP domains are colored as in Fig. 1. Created with BioRender.com.

Our simulations of closed mGBP2 provide a molecular picture of the initial docking step that precedes activation. Closed mGBP2 associated with all three membranes tested, in-

cluding the PS- and PC-containing controls, inserting its geranylgeranyl anchor as readily into these as into the cardiolipin-containing membrane. This is expected: a hydrophobic prenyl anchor inserts favorably irrespective of head-group chemistry, and the coarse-grained representation does not resolve the finer differences between lipid head groups. What distinguishes cardiolipin in our data is therefore not how often mGBP2 binds, but how it binds: CL forms specific contacts with lysine and arginine residues and becomes locally enriched at the binding site, whereas POPS, present at the same fraction of 5%, interacts about four times more weakly and forms diffuse, largely non-specific contacts. This is consistent with lipid-strip and GUV experiments, in which mGBP2 accumulates on membranes containing cardiolipin, phosphatidic acid, and monophosphorylated phosphoinositides, but not on phosphatidylserine-containing membranes,^32^ since these assays report stable accumulation over minutes rather than the initial adsorption of a single protein that our simulations capture.

Initial docking is also not sufficient for recruitment in cells. mGBP2 variants deficient in GTP binding (K51A) or isoprenylation (C586A) fail to localize to the *T. gondii* parasitophorous vacuole membrane, whereas the hydrolysis-deficient R48A variant still does,^59^ although the corresponding hGBP1 variant neither dimerizes nor disrupts membranes in vitro.^23,30^ Together with our finding that the closed dimer can insert only one of its two prenyl anchors and does not open (**Fig. 7A**), this suggests a two-stage recognition: prenyl-driven, largely unselective attachment of the closed protein, followed by GTP-dependent opening and oligomerization that converts transient attachment into stable accumulation. Lipid specificity would then act less as an on/off switch than as a bias on residence time.

A second finding concerns the conformational states that open GBPs adopt at membranes. In solution, the stability of the open state increased with oligomerization (**Fig. 7B-D**), from the monomer through the dimer to the hexamer, extending to mGBP2 what we previously found for hGBP1 and hGBP5^28^ and paralleling our simulations of the bacterial dynamin-like protein BDLP, which likewise requires higher-order assembly to maintain its open form.^65^ That this hinge motion is functionally relevant is supported by cross-linking experiments in which locking *α*12 to *α*7 abolished coat formation on bacteria, which was fully restored upon reduction.^30^ In our simulations, open monomers of both species folded back at membranes even though the GD was restrained to its nucleotide-bound conformation, whereas dimers and larger assemblies remained extended. The open state thus appears to be intrinsically unstable at the monomer level and to be maintained by oligomerization rather than by the nucleotide alone, consistent with the requirement for continuous GTP hydrolysis during coat assembly. ^30^

In our simulations, the two proteins folded back into different states. In hGBP1 on POPC, *α*12 came to rest on the side of the GD opposite to that of the crystal structures, contacting *α*0, with *α*12 and *α*13 forming one continuous helix. This arrangement corresponds to the conformer M2 described for hGBP1 in solution, which exchanges with the crystal-like conformer M1 on a 10–150 *µ*s timescale independently of bound nucleotide,^24^ and which our membrane-bound structure matches to within the resolution of the deposited ensemble. In contrast, mGBP2 on the PI5P membrane folded back with *α*12 on the crystal-structure side of the GD, contacting *α*3/4, and thus resembles M1. In both cases the prenyl anchor remained inserted in the bilayer rather than returning to its hydrophobic pocket, so these are membrane-bound resting states that the solution structures cannot capture (**Fig. 7A**). Whether the difference in *α*12 placement reflects an intrinsic preference of each protein, a directing role of the membrane, or the limited sampling of single trajectories cannot be decided from our data, as the two states formed on different membranes and mGBP2 did not fully close on POPC within 10 *µ*s. Notably, only a dimer formed from one M1 and one M2 protomer places the two *α*13 helices on the same side, which has been proposed as the prerequisite for the subsequent opening. ^24^ That both conformers arise spontaneously in our membrane simulations suggests that the fold-back states are not dead ends but the conformers from which a new open dimer can be built.

Our simulations suggest that neither oligomerization nor the bacterial outer membrane alone keeps open hGBP1 upright, but that the two act together. In solution, the stability of the open state increases from monomer to dimer to hexamer. On membranes, however, open monomers fold back, and open dimers, while remaining open, tip over unless they are supported. The hexamer stays intact on short-chain LPS but still tips over, whereas a single dimer remains upright on LPS with long O-antigen chains that reach up into the MiD region. In a laterally confined coat, in turn, all hGBP1 molecules remain upright even on POPC, because the bulky GDs and MiDs of neighboring molecules support each other, and only the void between the thin EDs near the membrane remains. This void lies at the height reached by short O-antigen chains. The O-antigen length required to keep open hGBP1 upright thus appears to depend on the degree of assembly: an isolated dimer needs long chains, whereas a laterally confined coat may need only short chains that fill the space between the EDs, or none at all. Lateral confinement and O-antigen length may therefore act complementarily (**Fig. 7E**).

This model is consistent with experimental observations that the O-antigen is dispensable for hGBP1 binding but matters for coat formation. hGBP1 binds both rough and smooth purified LPS,^66^ and bacteria with truncated O-antigen and outer core are coated and trigger downstream signaling.^30^ In our simulations, membrane anchoring is likewise mediated by the farnesyl anchor and the adjacent polybasic motif, independent of the O-antigen. In the dense native coat, where roughly 12,000 hGBP1 molecules cover a single minicell, ^30^ lateral confinement is complete, and our coat simulation shows that this alone suffices to keep open hGBP1 upright. Where confinement is incomplete, however, the O-antigen becomes relevant: hGBP1 binds intact *S. flexneri* more efficiently when the O-antigen is present, and the transition from bacteria-attached polymers to a uniform, bacteria-encasing coat depends on O-antigen chains;^66^ coat formation is moreover cooperative and sets in above a threshold concentration.^23,30^ Our simulations offer a structural explanation: at coat edges and during nucleation, where neighbors are missing, long O-antigen chains could hold individual hGBP1 dimers upright until lateral support is established. This leads to a testable prediction: O-antigen length should matter most in the early stages of coat formation and least within a mature coat.

Although the two species target different membranes, our simulations suggest that they face the same problem. On the PI5P membrane, the open mGBP2 dimer lies flat rather than standing upright, but it remains open: the hinge at the MiD–ED transition does not close, and only the C-terminus kinks, held down by strong interactions between PI5P and the protein. In fact, none of the open dimers closed at this hinge on any membrane; what differs is the flexibility of the EDs, which is greatest for hGBP1 on LPS, where the O-antigen chains pull on the stalks. The mGBP2 dimer is therefore not inactivated by lying down, and lateral confinement within a coat should be able to raise it, as we observed for hGBP1 on a plain POPC membrane. Coat formation of the kind resolved for hGBP1^23,30^ has not been demonstrated for mGBP2, which forms supramolecular complexes at the *T. gondii* parasitophorous vacuole membrane, ^59^ but our results predict that it would resolve the same problem in the same way. What differs between the species is thus not the mechanism but the lipid that identifies the target: in both cases the characteristic lipid of the membrane, LPS for hGBP1 and PI5P for mGBP2, accounts for most of the protein–lipid interaction, whereas the remaining lipids interact mainly with the C-terminal anchor region. Which lipid code allows mGBP2 to discriminate the vacuolar membrane from host membranes remains unresolved. PI5P has been proposed as such a signal for the immunity-related GTPase Irgb6, which co-localizes with mGBPs at the parasitophorous vacuole membrane and preferentially binds phosphoinositides and phosphatidylserine; ^53^ our simulations indicate that phosphoinositide-containing patches likewise suffice to anchor mGBP2 through head-group interactions.

The disc of 30 hGBP1 dimers (**Fig. 7D**), finally, speaks to the proposed function of the polymeric form as a protein depot from which GBP1 is delivered to membranes.^45,66^ In our simulation the disc separated into smaller clusters, consistent with the observation that the lipid-free polymeric form is micellar rather than disc-shaped, ^23^ while individual farnesyl anchors inserted into the bilayer. Direct insertion from the polymeric form thus appears feasible, whereas the disc itself left the membrane largely undisturbed, indicating that membrane damage arises from the assembled coat rather than from contact with the polymer.

Several limitations should be kept in mind. Owing to system sizes of up to 7.5 million beads, most open-state and oligomer simulations were performed as single trajectories of 1– 10 *µ*s. We therefore base our conclusions not on individual events but on patterns that recur across independent systems: all open monomers folded back or kinked, whereas all open dimers remained extended, for both species and on all membranes tested, mirroring their relative stability in solution. Within the oligomers, the individual dimers provide internal replicates, and the long-chain LPS dimer, the only system simulated twice, reproduced the same behavior. Statements about single events, such as the initial docking of the disc, should be regarded as tendencies.

The comparison between short- and long-chain O-antigen is further confounded by differences in resolution, force field, and oligomeric state, as the short-chain systems were simulated coarse-grained with MARTINI 2.2 and the long-chain dimer all-atom. MAR-TINI 2 overestimates protein–protein attraction, which was reduced in MARTINI 3,^39,62^ so the greater cohesion of the hexamer on LPS relative to the hexamer on POPC may partly reflect the force field; the tipping itself, however, was also observed for monomers and dimers on POPC, where no sugars are present. Finally, no nucleotide was present in the coarse-grained simulations, so nucleotide-dependent dynamics are not captured, although the GD was restrained to its nucleotide-bound conformation; the coarse-grained LPS model offers only short O-antigen chains; and the 1–10 *µ*s timescale samples the initial phase of membrane engagement rather than coat assembly, which proceeds over minutes in cells.^30^

## 4 Conclusion

Our simulations describe how GBPs engage pathogen membranes between the two states that structural experiments resolve: the closed protein in solution and the assembled coat. Closed GBPs dock through their prenyl anchor with little lipid selectivity, and selectivity instead resides in the quality of the subsequent contacts. Once opened, however, the extended protein is intrinsically unstable on its own: monomers fold back into membrane-bound resting states, and dimers tip over unless they are held. Two things can hold them, the O-antigen layer of the bacterial outer membrane and the lateral support of neighboring molecules within a coat, and long O-antigen chains can substitute for support from neighbors. The same should apply to murine GBP2 on phosphoinositide-containing membranes, where the open dimer lies flat but remains open. This places the decisive step at the edge of a growing coat, where a GBP must stay extended before its neighbors are in place. This regime is hard to reach experimentally but well suited to simulation, and is the natural target for future work, alongside structural characterization of extended mGBP2.

## 5 Methods

### 5.1 Preparation of the GBP structures

#### CG models and membrane with closed mGBP2

The CG models of the proteins were created with the Python script *martinize.py*,^37^ and the membranes were built with *insane.py* .^38^ The secondary structure was provided as an input file. The membranes were equilibrated for 1 *µ*s before the protein was added, and mGBP2 was placed 4 nm above the membrane surface using PyMOL^35^ so that no protein–membrane contacts were present at the start of the simulations. After solvation and neutralization, water beads trapped between the two leaflets were removed with PyMOL.

#### CG models of open hGBP1 and mGBP2 and membranes

The CG models of the open proteins and the LPS membranes were created with CHARMM-GUI,^36^ which converts atomistic PDB input to the CG level and places the protein in the membrane. The protein– LPS membrane systems were solvated and neutralized with CaCl_2_ for the LPS core, with an additional 150 mM NaCl. Since the open proteins are anchored in the membrane from the start, no preceding equilibration of the protein above the membrane was required. These systems were simulated with MARTINI 2.2, the only version of MARTINI in which LPS is available. All other open-state systems were built with the *martinize.py* programme^41^ and *insane.py* ^38^ and simulated with MARTINI 3, which provides parameters for phosphoinositides and prenyl groups and an improved description of protein–protein interactions. Solvation, neutralization, minimization, equilibration, and production followed the same schemes as for the LPS systems.

#### CG models of the coat-like and disc assemblies of hGBP1 with POPC membranes

The coat-like hGBP1 model was constructed from three hexamers, i.e. 18 hGBP1 molecules, based on the PDB structure 8R1A, arranged such that the hexamers touch their periodic images on a 20 *×* 27 nm^2^ POPC membrane. This created an effectively infinite protein layer in the membrane plane and restricted the lateral movement of the cone-shaped hGBP1 molecules. The disc, comprising 30 hGBP1 dimers (60 molecules), was built after the cryo-EM model of the polymeric assembly of hGBP1 (EMD-18698).^22^ We aligned the individual hGBP1 dimer units to the corresponding cryo-EM density in ChimeraX,^67^ moving them 2.5 nm outwards to avoid steric clashes between the stalk residues at the resolution of the map. CG models of both assemblies were generated with *martinize2*,^41^ and the POPC membranes were built with *insane.py* ^38^ as for the other MARTINI 3 systems. The disc was placed about 4 nm above a 100 *×* 100 nm^2^ POPC membrane. The systems were solvated, ionized with 150 mM NaCl, and equilibrated following the same minimization and equilibration scheme as the other MARTINI 3 membrane systems.

### 5.2 Simulation details

#### CGMD simulations of the closed GBP systems

The energy of the closed mGBP2 membrane systems was first minimized using the steepest descent algorithm, followed by equilibration to the desired temperature of 310 K and pressure of 1 bar to mimic the physiological environment. For a slower equilibration, a 0.25 ns *NpT* equilibration with a time step of 10 fs was applied followed by a 0.5 ns *NpT* equilibration with a time step of 20 fs. During these steps, the protein’s backbone beads were restrained, allowing the water molecules to relax around the solute. The velocity rescaling thermostat^68^ and Berendsen barostat^69^ were employed to regulate the temperature and pressure in the *NpT* simulations (10 fs time step), while the semi–isotropic Parrinello-Rahman barostat^70^ was used for the *NpT* simulations with 20 fs time step. The PME sum method^71,72^ was used to calculate long-range electrostatic interactions with periodic boundary conditions, while ring systems and stiff bonds constraints were applied via the LINCS algorithm.^73^ The vdW and Coulombic interaction cutoffs were set to 1.2 nm. Production MD runs were performed for 1-10 *µ*s for each system and the time step was set to 20 fs for systems, where the coordinates and velocities were recorded every 100 ps.

#### CGMD simulations of the open GBP systems

The start structures were obtained from our previous work,^28^ from AlphaFold, the PDB data bank (PDB: 8CQB and 8R1A)^22,23^ and from the homology modeling software SwissModel. Some structures needed the extension of EDs manually which we did with PyMOL. The prenyl groups were also added at the EDs using PyMOL. The mGBP2 sequence was fitted onto the PDB template of the hGBP1 dimer (PDB: 8CQB) via SwissModel to obtain a similar criss-cross protein structure that provides better stability. Like in our previous work the CG parameters of the prenyl groups for all MARTINI 2 runs were taken from the work by Atsmon-Raz et al. ^74^. Additionally, the prenyl parameters for the MARTINI 3 runs were taken from Koukos et al., ^40^ built via CHARMM-GUI’s PDB Reader & Manipulator, and translated into CG coordinates via the martinize2 programme.^41^

Depending on the system (see Table 1), the simulations were performed with the MAR-TINI 2.2^37,75^ or the MARTINI 3^39^ force field. For the elastic network ElNeDyn applied in MARTINI 2, the constraints between the beads were built for distances between 0.5 and 0.9 nm with a force of 500 kJ mol^-1^ nm^-2^ ^76^. Furthermore, we used the domELNEDIN method^77^ to remove the constraints between the domains, which was applied as a Tcl script within the Visual Molecular Dynamics (VMD) program^78^. For MARTINI 3 we applied the basic elastic network model and commented out the interdomain rubber bands to match the elastic network method domELNEDIN from above, as this is compatible with the new parameter and implementation set in the martinize2 programme. The parameters for LPS were taken from the Martini Maker of CHARMM-GUI,^36,79,80^ using the variant with the longest available O-antigen (4 repeat units). For the AAMD simulations, LPS with 12 O-antigen repeat units was used, matching the average chain length of *S. flexneri* .^61^ The parameters for the phosphoinositide lipids were taken from Borges-Araújo et al. ^81^

CHARMM-GUI provides GROMACS compatible input files, including the force fields, a solvated PDB structure, and ready-to-use simulation inputs.^36^ These inputs include two energy minimization steps using soft-core potentials and the steepest descent algorithm to eliminate bead clashes, followed by five semi-isobaric *NpT* equilibration steps at 310 K and 1 bar, each applying gradually reduced positional restraints on the system. The production MD run with an *NpT* ensemble, applies the Nose–Hoover^82,83^ thermostat and the Parrinello-Rahman barostat.^70^ Periodic boundary conditions are employed throughout. Non-bonded interactions are treated with 1.2 nm cutoffs for both vdW and Coulombic interactions, and long-range electrostatics are calculated using the reaction field method for faster performance. Production MD runs were performed for 1-10 *µ*s for each system and the time step was set to 20 fs, where the coordinates and velocities were recorded every 100 ps. The same protocol was used for the open-state simulations in solution, except that isotropic pressure coupling was applied and the simulation time was limited to 5 *µ*s per run.

#### AAMD simulations

The AAMD simulations of the hGBP1 dimer with the long O-antigen LPS membrane were set up using CHARMM-GUI’s atomistic membrane builder and their GROMACS compatible input files. ^36^ Farnesylation and positioning in the membrane was directly done in CHARMM-GUI. The protein–LPS membrane systems were solvated, and neutralized with CaCl_2_ for the LPS core and additional 150 mM NaCl. We used the CHARMM-GUI minimization, equilibration and production schemes to run the simulations with the C36m force field.^84^ This includes two energy minimization using the steepest descent algorithm to eliminate atom clashes, followed by five semi-isobaric *NpT* equilibration steps at 310 K and 1 bar, each applying gradually reduced positional restraints on the system. The semiisotropic production MD run with an *NpT* ensemble, applying the Nose–Hoover^82,83^ thermostat and the Parrinello-Rahman barostat.^70^ The runs were performed for 1*µ*s with a time step of 2 fs. Periodic boundary conditions are employed throughout. Non-bonded interactions are treated with 1.2 nm cutoffs for both vdW and Coulombic interactions, and long-range electrostatics are calculated using the PME method.

### 5.3 Analysis

#### Density plots

To check whether lipids clustered around the protein in the *xy*-plane, we generated density maps of GBPs and the lipids via *gmx densmap*. Here, the direction to average was over *z*-axis, the grid size was set to 0.08 and the unit was nm^-3^. We changed the format from xpm to a matrix dat file via the DulvyTools^85^ (*dit xpm2dat*). Subsequently, this could be used for plotting the density maps via an own python script. To standardize the axis and colors, we selected the range between 0 and 1.5 nm^-3^ for DOPC in yellow and CHOL in blue, between 0 and 0.2 nm^-3^ for CL in magenta and between 0 and 0.4 nm^-3^ for GBPs in green. We additionally used yellow for the extended proteins, purple for POPC, red for LPS, blue for PI5P, gray for POPE, green for POPG, and orange for CL.

#### Δ*z*-position analysis

We observed the motions of the three mGBP2 domains regarding the membranes by calculating the distance of the centers of mass of the three domains concerning the membrane surface along the *z* coordinate. To define the membrane surface position, the average position of the PO4 lipid beads were used. To further specify which parts of the protein are closest to the membrane, we also calculated the distances from the membrane surface of the GD, of residue L480 at the MiD–ED transition, which forms the tip of the closed protein opposite the GD, and of the geranylgeranyl anchor. This analysis was conducted for every 1 ns.

#### Lipid number analysis

To study the GBP–lipid interactions, we calculated the time-averaged and cumulative lipid numbers over the entire simulations and over the binding events, using the lipid fingerprint script provided by the MARTINI developers.^86^ The normalized lipid number was obtained by dividing the average lipid number by the fraction of that lipid in the membrane (DOPC: 1140 of 1520 lipids, or 1216 for PC-mem; CHOL: 304; CL or POPS: 76). A distance cut-off of 1.0 nm was applied, using the same protein parts as in the Δ*z*-position analysis. For each binding pose and membrane composition, we further calculated the weighted mean and standard deviation, weighting each simulation by the number of frames spent in that pose. The analysis was based on 1000 frames per simulation.

#### Protein–lipid interaction

We analyzed the protein–lipid interactions with the PyLipID package of Song et al.,^60^ which measures lipid residence times, generates representative lipid poses, and distinguishes unbinding events from transient interruptions of contact. For the latter, a dual-cutoff scheme is used, in which a contact starts when the distance falls below the lower cut-off and ends only when it exceeds the upper one; we used the default values of 0.475 and 0.7 nm. We analyzed every 10th frame of the trajectories after correction for periodic boundary conditions, specifying the number of protein chains and an all-atom PDB file to map the binding sites onto the structure. The three residues with the highest lipid occupancy, excluding the lipid anchors, are reported.

#### Energy analysis

We calculated interaction energies (*E_int_*) between distinct lipid components and each protein residue. Primarily, to determine whether electrostatic or van der Waals (vdW) forces dominate protein-lipid membrane interactions. The total energy (*E_total_*) comprises short-range Coulomb (*E_Coul_*) and Lennard-Jones (*E_LJ_*) potentials. We focused on significant interactions with absolute values greater than 2 kJ/mol for each protein-lipid pair. Secondly, we categorized the residues into five groups for *E_int_*: 1. Polar (Ser, Thr, Asn, Gln) 2. Nonpolar (Gly, Ala, Val, Leu, Ile, Pro, Met, Cys) 3. Aromatic (Phe, yr, Trp) 4. Negative (Asp, Glu) 5. Positive (Lys, Arg, His). The interaction energies were normalized by the number of residues in contact with each lipid component, counting only residues with a non-zero interaction energy (*|E| >* 10*^−^*^6^ kJ mol*^−^*^1^).

#### Backmapping procedure

To translate the CG-level GBP models back to the AA level, we used the *Backward* method.^87,88^ Here, we project the CG coordinates to AA coordinates via *backward.py*, correct the coordinates with *initram.sh*, and relax the final structures using energy minimization in vacuum.

## Acknowledgement

B.S. gratefully acknowledges funding from the Deutsche Forschungsgemeinschaft (DFG, German Research Foundation) (project A07 of the CRC 1208). The project was made possible thanks to computing time granted through the Gauss Centre for Supercomputing e.V. (www.gauss-centre.eu) on the GCS Supercomputer SuperMUC-NG at Leibniz Supercomputing Centre (www.lrz.de) (project pn98zo). Computational infrastructure and support were provided by the Centre for Information and Media Technology at Heinrich Heine University Düsseldorf.

## References

(1) MacMicking, J. D. Interferon-inducible effector mechanisms in cell-autonomous immunity. Nat. Rev. Immunol. 2012, 12, 367–382.

(2) Ghosh, A.; Praefcke, G. J. K.; Renault, L.; Wittinghofer, A.; Herrmann, C. How guanylate-binding proteins achieve assembly-stimulated processive cleavage of GTP to GMP. Nature 2006, 440, 101–104.

(3) Lorenz, C.; Ince, S.; Zhang, T.; Cousin, A.; Batra-Safferling, R.; Nagel-Steger, L.; Herrmann, C.; Stadler, A. M. Farnesylation of human guanylate-binding protein 1 as safety mechanism preventing structural rearrangements and uninduced dimerization. FEBS J. 2019, 287, 496–514.

(4) Tretina, K.; Park, E.-S.; Maminska, A.; MacMicking, J. D. Interferon-induced guanylate-binding proteins: Guardians of host defense in health and disease. J. Exp. Med. 2019, 216, 482–500.

(5) Sistemich, L.; Kutsch, M.; Hämisch, B.; Zhang, P.; Shydlovskyi, S.; Britzen-Laurent, N.; Stürzl, M.; Huber, K.; Herrmann, C. The Molecular Mechanism of Polymer Formation of Farnesylated Human Guanylate-binding Protein 1. J. Mol. Biol. 2020, 432, 2164–2185.

(6) Britzen-Laurent, N.; Bauer, M.; Berton, V.; Fischer, N.; Syguda, A.; Reipschläger, S.; Naschberger, E.; Herrmann, C.; Stürzl, M. Intracellular Trafficking of Guanylate-Binding Proteins Is Regulated by Heterodimerization in a Hierarchical Manner. PLoS One 2010, 5, e14246.

(7) Legewie, L.; Loschwitz, J.; Steffens, N.; Prescher, M.; Wang, X.; Smits, S. H.; Schmitt, L.; Strodel, B.; Degrandi, D.; Pfeffer, K. Biochemical and structural characterization of murine GBP7, a guanylate binding protein with an elongated C-terminal tail. Biochem. J. 2019, 476, 3161–3182.

(8) Cui, W.; Wang, T.; Feng, J. From mechanisms to therapies: the multifaceted roles of guanylate-binding protein 2 in immunity, cancer, and beyond. Front. Immunol. 2025, 16.

(9) Shi, Y.; Chichung Lie, D.; Taupin, P.; Nakashima, K.; Ray, J.; Yu, R. T.; Gage, F. H.; Evans, R. M. Expression and function of orphan nuclear receptor TLX in adult neural stem cells. Nature 2004, 427, 78–83.

(10) Yamamoto, M.; Okuyama, M.; Ma, J. S.; Kimura, T.; Kamiyama, N.; Saiga, H.; Ohshima, J.; Sasai, M.; Kayama, H.; Okamoto, T.; Huang, D. C.; Soldati-Favre, D.; Horie, K.; Takeda, J.; Takeda, K. A Cluster of Interferon-*γ*-Inducible p65 GTPases Plays a Critical Role in Host Defense against Toxoplasma gondii. Immunity 2012, 37, 302–313.

(11) Man, S. M.; Karki, R.; Malireddi, R. K. S.; Neale, G.; Vogel, P.; Yamamoto, M.; Lamkanfi, M.; Kanneganti, T.-D. The transcription factor IRF1 and guanylate-binding proteins target activation of the AIM2 inflammasome by Francisella infection. Nat. Immunol. 2015, 16, 467–475.

(12) Santos, J. C.; Boucher, D.; Schneider, L. K.; Demarco, B.; Dilucca, M.; Shkarina, K.; Heilig, R.; Chen, K. W.; Lim, R. Y. H.; Broz, P. Human GBP1 binds LPS to initiate assembly of a caspase-4 activating platform on cytosolic bacteria. Nat. Commun. 2020, 11.

(13) Kim, B.-H.; Shenoy, A. R.; Kumar, P.; Das, R.; Tiwari, S.; MacMicking, J. D. A Family of IFN-*γ*–Inducible 65-kD GTPases Protects Against Bacterial Infection. Science 2011, 332, 717–721.

(14) Jacot, D.; Frenal, K.; Marq, J.-B.; Sharma, P.; Soldati-Favre, D. Assessment of phospho-rylation inToxoplasmaglideosome assembly and function: Phosphorylation and glideosome function. Cell. Microbiol. 2014, 16, 1518–1532.

(15) Olszewski, M. A.; Gray, J.; Vestal, D. J. In SilicoGenomic Analysis of the Human and Murine Guanylate-Binding Protein (GBP) Gene Clusters. J. Interferon Cytokine Res. 2006, 26, 328–352.

(16) Kutsch, M.; Coers, J. Human guanylate binding proteins: nanomachines orchestrating host defense. FEBS J. 2021, 288, 5826–5849.

(17) Kohler, K. M.; Kutsch, M.; Piro, A. S.; Wallace, G. D.; Coers, J.; Barber, M. F. A Rapidly Evolving Polybasic Motif Modulates Bacterial Detection by Guanylate Binding Proteins. mBio 2020, 11.

(18) Degrandi, D.; Kravets, E.; Konermann, C.; Beuter-Gunia, C.; Klümpers, V.; Lahme, S.; Wischmann, E.; Mausberg, A. K.; Beer-Hammer, S.; Pfeffer, K. Murine Guanylate Binding Protein 2 (mGBP2) controls Toxoplasma gondii replication. Proc. Natl. Acad. Sci. U. S. A. 2013, 110, 294–299.

(19) Ĉorte-Real, J. V.; Baldauf, H.-M.; Melo-Ferreira, J.; Abrantes, J.; Esteves, P. J. Evolution of Guanylate Binding Protein (GBP) Genes in Muroid Rodents (Muridae and Cricetidae) Reveals an Outstanding Pattern of Gain and Loss. Front. Immunol. 2022, 13.

(20) Schelle, L.; Corte-Real, J. V.; Esteves, P. J.; Abrantes, J.; Baldauf, H.-M. Functional cross-species conservation of guanylate-binding proteins in innate immunity. Med. Microbiol. Immunol. 2022, 212, 141–152.

(21) Neuweiler, H. Decision letter: Integrative dynamic structural biology unveils conformers essential for the oligomerization of a large GTPase. 2022; 10.7554/elife.79565.sa1.

(22) Weismehl, M.; Chu, X.; Kutsch, M.; Lauterjung, P.; Herrmann, C.; Kudryashev, M.; Daumke, O. Structural insights into the activation mechanism of antimicrobial GBP1. EMBO J. 2024, 43, 615–636.

(23) Kuhm, T.; Taisne, C.; de Agrela Pinto, C.; Gross, L.; Giannopoulou, E. A.; Huber, S. T.; Pardon, E.; Steyaert, J.; Tans, S. J.; Jakobi, A. J. Structural basis of antimicrobial membrane coat assembly by human GBP1. Nat. Struct. Mol. Biol. 2024, 32, 172–184.

(24) Peulen, T.-O. et al. Integrative dynamic structural biology unveils conformers essential for the oligomerization of a large GTPase. eLife 2023, 12.

(25) Cui, W. et al. Structural basis for GTP-induced dimerization and antiviral function of guanylate-binding proteins. Proc. Natl. Acad. Sci. U. S. A. 2021, 118.

(26) Roy, S.; Wang, B.; Tian, Y.; Yin, Q. Crystal structures reveal nucleotide-induced conformational changes in G motifs and distal regions in guanylate-binding protein 2. *bioRxiv* 2023, Preprint.

(27) Roy, S.; Wang, B.; Roy, K.; Tian, Y.; Bhattacharya, M.; Williams, S.; Yin, Q. Crystal structures reveal nucleotide-induced conformational changes in G motifs and distal regions in human guanylate-binding protein 2. *Commun*. Biol. 2025, 8.

(28) Bundschuh, B. F.; Loschwitz, J.; Strodel, B. Mechanisms driving open-closed transitions and dimer stability of the large GTPases hGBP1 and hGBP5. Front. Mol. Biosci. 2026, 13.

(29) Kutsch, M.; Herrmann, C. Glue it together: self-assembly of the antibacterial GTPase GBP1. Biol. Chem. 2026, 407, 105–122.

(30) Zhu, S.; Bradfield, C. J.; Maminska, A.; Park, E.-S.; Kim, B.-H.; Kumar, P.; Huang, S.; Kim, M.; Zhang, Y.; Bewersdorf, J.; MacMicking, J. D. Native architecture of a human GBP1 defense complex for cell-autonomous immunity to infection. Science 2024, 383.

(31) Abramson, J. et al. Accurate structure prediction of biomolecular interactions with AlphaFold 3. Nature 2024, 630, 493–500.

(32) Loschwitz, J.; Steffens, N.; Wang, X.; Schäffler, M.; Pfeffer, K.; Degrandi, D.; Strodel, B. Domain motions, dimerization, and membrane interactions of the murine guanylate binding protein 2. Sci. Rep. 2023, 13, 1–20.

(33) Schumann, W.; Loschwitz, J.; Reiners, J.; Degrandi, D.; Legewie, L.; Stühler, K.; Pfeffer, K.; Poschmann, G.; Smits, S. H. J.; Strodel, B. Integrative modeling of guanylate binding protein dimers. Protein Sci. 2023, 32.

(34) Waterhouse, A.; Bertoni, M.; Bienert, S.; Studer, G.; Tauriello, G.; Gumienny, R.; Heer, F. T.; de Beer, T. A. P.; Rempfer, C.; Bordoli, L.; Lepore, R.; Schwede, T. SWISS-MODEL: homology modelling of protein structures and complexes. Nucleic Acids Res. 2018, 46, W296–W303.

(35) Schrodinger, LLC The PyMOL Molecular Graphics System, Version 3.0. 2025; https://pymol.org.

(36) Jo, S.; Kim, T.; Iyer, V. G.; Im, W. CHARMM-GUI: a web-based graphical user interface for CHARMM. J. Comput. Chem. 2008, 29, 1859–1865.

(37) de Jong, D. H.; Singh, G.; Bennett, W. F. D.; Arnarez, C.; Wassenaar, T. A.; Schäfer, L. V.; Periole, X.; Tieleman, D. P.; Marrink, S. J. Improved Parameters for the Martini Coarse-Grained Protein Force Field. J. Chem. Theory Comput. 2013, 9, 687–697.

(38) Wassenaar, T. A.; Ingolfsson, H. I.; Bockmann, R. A.; Tieleman, D. P.; Marrink, S. J. Computational Lipidomics with insane: A Versatile Tool for Generating Custom Membranes for Molecular Simulations. J. Chem. Theory Comput. 2015, 11, 2144–2155, PMID: 26574417.

(39) Souza, P. C.; Alessandri, R.; Barnoud, J.; Thallmair, S.; Faustino, I.; Grünewald, F.; Patmanidis, I.; Abdizadeh, H.; Bruininks, B. M.; Wassenaar, T. A.; others Martini 3: a general purpose force field for coarse-grained molecular dynamics. Nat. Methods 2021, 18, 382–388.

(40) Koukos, P. I.; Dehghani-Ghahnaviyeh, S.; Velez-Vega, C.; Manchester, J.; Tieleman, D. P.; Duca, J. S.; Souza, P. C. T.; Cournia, Z. Martini 3 Force Field Parameters for Protein Lipidation Post-Translational Modifications. J. Chem. Theory Comput. 2023, 19, 8901–8918.

(41) Kroon, P. C.; Grunewald, F.; Barnoud, J.; van Tilburg, M.; Brasnett, C.; de Souza, P. C. T.; Wassenaar, T. A.; Marrink, S.-J. J. Martinize2 and Vermouth: Unified Framework for Topology Generation. eLife 2025, 13.

(42) Huang, J.; Rauscher, S.; Nawrocki, G.; Ran, T.; Feig, M.; de Groot, B.; Grubmüller, H.; MacKerell Jr, A. CHARMM36m: an improved force field for folded and intrinsically disordered proteins. Nat. Methods 2017, 14, 71.

(43) Steffens, N. Analyse der dynamischen mGBP-Rekrutierung zu Pathogen-enthaltenden Membrankompartimenten. PhD thesis, Heinrich-Heine-Universität, Düsseldorf, Deutschland, 2020.

(44) Steffens, N.; Beuter-Gunia, C.; Kravets, E.; Reich, A.; Legewie, L.; Pfeffer, K.; Degrandi, D.; Sibley, L. D. Essential Role of mGBP7 for Survival of Toxoplasma gondii Infection. mBio 2020, 11, e02993–19.

(45) Shydlovskyi, S.; Zienert, A.; Ince, S.; Dovengerds, C.; Hohendahl, A.; Dargazanli, J.; Blum, A.; Günther, S.; Kladt, N.; Stürzl, M.; Schauss, A.; Kutsch, M.; Roux, A.; Praefcke, G.; Herrmann, C. Nucleotide-dependent farnesyl switch orchestrates polymerization and membrane binding of human guanylate-binding protein 1. Proc. Natl. Acad. Sci. U. S. A. 2017, 114, E5559–E5568.

(46) Rana, F. R.; Sultany, C. M.; Blazyk, J. Determination of the lipid composition of Salmonella typhimurium outer membranes by 31P NMR. J. Microbiol. Methods 1991, 14, 41–51.

(47) Nikaido, H. Molecular Basis of Bacterial Outer Membrane Permeability Revisited. Microbiol. Mol. Biol. Rev. 2003, 67, 593–656.

(48) Horne, J. E.; Brockwell, D. J.; Radford, S. E. Role of the lipid bilayer in outer membrane protein folding in Gram-negative bacteria. J. Biol. Chem. 2020, 295, 10340–10367.

(49) Brahmbhatt, H. N.; Lindberg, A. A.; Timmis, K. N. Pathogenesis of Shigellosis; Springer Berlin Heidelberg, 1992; pp 45–64.

(50) Hancock, R. E.; Karunaratne, D. N.; Bernegger-Egli, C. In Bacterial Cell Wall ; Ghuysen, J.-M., Hakenbeck, R., Eds.; New Comprehensive Biochemistry; Elsevier, 1994; Vol. 27; pp 263–279.

(51) Welti, R.; Mui, E.; Sparks, A.; Wernimont, S.; Isaac, G.; Kirisits, M.; Roth, M.; Roberts, C. W.; Botte, C.; Marechal, E.; McLeod, R. Lipidomic Analysis of ¡i¿Toxoplasma gondii¡/i¿ Reveals Unusual Polar Lipids. Biochemistry 2007, 46, 13882– 13890.

(52) Besteiro, S.; Bertrand-Michel, J.; Lebrun, M.; Vial, H.; Dubremetz, J.-F. Lipidomic analysis of Toxoplasma gondii tachyzoites rhoptries: further insights into the role of cholesterol. Biochem. J. 2008, 415, 87–96.

(53) Lee, Y.; Yamada, H.; Pradipta, A.; Ma, J. S.; Okamoto, M.; Nagaoka, H.; Takashima, E.; Standley, D. M.; Sasai, M.; Takei, K.; Yamamoto, M. Initial phospholipid-dependent Irgb6 targeting to¡i¿Toxoplasma gondii¡/i¿vacuoles mediates host defense. Life Sci. Alliance 2019, 3, e201900549.

(54) Cruz-Bustos, T.; Joachim, A. Toxoplasma gondii assembles extracellular vesicles with conserved lipid profiles across host cell types. Front. Cell. Infect. Microbiol. 2026, 16.

(55) Prakash, B.; Renault, L.; Praefcke, G.; Herrmann, C.; Wittinghofer, A. Triphosphate structure of guanylate-binding protein 1 and implications for nucleotide binding and GTPase mechanism. EMBO J. 2000, 19, 4555–4564.

(56) Ji, C.; Du, S.; Li, P.; Zhu, Q.; Yang, X.; Long, C.; Yu, J.; Shao, F.; Xiao, J. Structural mechanism for guanylate-binding proteins (GBPs) targeting by the Shigella E3 ligase IpaH9.8. PLoS Pathog. 2019, 15, e1007876.

(57) Kravets, E.; Degrandi, D.; Weidtkamp-Peters, S.; Ries, B.; Konermann, C.; Felekyan, S.; Dargazanli, J. M.; Praefcke, G. J.; Seidel, C. A.; Schmitt, L.; others The GTPase activity of murine guanylate-binding protein 2 (mGBP2) controls the intracellular localization and recruitment to the parasitophorous vacuole of *Toxoplasma gondii*. J. Biol. Chem. 2012, 287, 27452–27466.

(58) Barz, B.; Loschwitz, J.; Strodel, B. Large-scale, dynamin-like motions of the human guanylate binding protein 1 revealed by multi-resolution simulations. PLoS Comput. Biol. 2019, 15, 1–29.

(59) Kravets, E.; Degrandi, D.; Ma, Q.; Peulen, T.-O.; Klümpers, V.; Felekyan, S.; Kühnemuth, R.; Weidtkamp-Peters, S.; Seidel, C. A.; Pfeffer, K. Guanylate binding proteins directly attack *Toxoplasma gondii* via supramolecular complexes. eLife 2016, 5, e11479.

(60) Song, W.; Corey, R. A.; Ansell, T. B.; Cassidy, C. K.; Horrell, M. R.; Duncan, A. L.; Stansfeld, P. J.; Sansom, M. S. P. PyLipID: A Python package for analysis of protein-lipid interactions from molecular dynamics simulations. J. Chem. Theory Comput. 2022, 18, 1188–1201.

(61) Hoare, A.; Bravo, D.; Martinic, M.; Valvano, M. A.; Contreras, I.; Alvarez, S. A. The normal chain length distribution of the O antigen is required for the interaction of Shigella flexneri 2a with polarized Caco-2 cells. Biol. Res. 2012, 45, 21–26.

(62) Javanainen, M.; Martinez-Seara, H.; Vattulainen, I. Excessive aggregation of membrane proteins in the Martini model. PLoS One 2017, 12, e0187936.

(63) Fuhrmans, M.; Müller, M. Coarse-grained simulation of dynamin-mediated fission. Soft Matter 2015, 11, 1464–1480.

(64) Mahmood, M. I.; Kubo, S.; Noguchi, H.; Okazaki, K.-i. Membrane Constriction by Dynamin through GTP-Driven Conformational Changes from Coarse-Grained Molecular Dynamics Simulations. J. Phys. Chem. Lett. 2025, 16, 12411–12418.

(65) Schumann, W.; Bundschuh, B. F.; Loschwitz, J.; Strodel, B. From Closed to Mighty: How GTP-Induced Allosteric Dynamics, Dimerization, and Membrane Binding Transform a Bacterial Dynamin-like Protein. Journal of Chemical Information and Modeling 2026,

(66) Kutsch, M.; Sistemich, L.; Lesser, C. F.; Goldberg, M. B.; Herrmann, C.; Coers, J. Direct binding of polymeric GBP1 to LPS disrupts bacterial cell envelope functions. EMBO J. 2020, 39, e104926.

(67) Meng, E. C.; Goddard, T. D.; Pettersen, E. F.; Couch, G. S.; Pearson, Z. J.; Morris, J. H.; Ferrin, T. E. ¡scp¿UCSF ChimeraX¡/scp¿: Tools for structure building and analysis. Protein Sci. 2023, 32.

(68) Bussi, G.; Donadio, D.; Parrinello, M. Canonical sampling through velocity rescaling. J. Chem. Phys. 2007, 126, 014101.

(69) Berendsen, H.; Postma, J.; van Gunsteren, W.; Di Nola, A.; Haak, J. Molecular dynamics with coupling to an external bath. Journal of Chemical Physics 1984, 81, 3684–3690.

(70) Parrinello, M.; Rahman, A. Polymorphic Transitions in Single-Crystals - a New Molecular-Dynamics Method. Mol. Phys. 1981, 52, 7182–7190.

(71) Darden, T.; York, D.; Pedersen, L. PARTICLE Mesh Ewald - an N.Log(N) Method for Ewald Sums in Large Systems. J. Chem. Phys. 1993, 98, 10089–10092.

(72) Essmann, U.; Perera, L.; Berkowitz, M. L. A Smooth PARTICLE Mesh Ewald Method. J. Chem. Phys. 1995, 103, 8577–8593.

(73) Hess, B.; Bekker, H.; Berendsen, H. J. C.; Fraaije, J. LINCS: A linear constraint solver for molecular simulations. J. Comput. Chem. 1997, 18, 1463–1472.

(74) Atsmon-Raz, Y.; Tieleman, D. P. Parameterization of Palmitoylated Cysteine, Farnesylated Cysteine, Geranylgeranylated Cysteine, and Myristoylated Glycine for the Martini Force Field. J. Phys. Chem. B 2017, 121, 11132–11143.

(75) Marrink, S.; Risselada, H.; Yefimov, S.; Tieleman, D.; de Vries, A. The MARTINI Force Field: Coarse Grained Model for Biomolecular Simulations. J. Phys. Chem. B 2007, 111, 7812–7824.

(76) Periole, X.; Cavalli, M.; Marrink, S.-J.; Ceruso, M. A. Combining an Elastic Network With a Coarse-Grained Molecular Force Field: Structure, Dynamics, and Intermolecular Recognition. J. Chem. Theory Comput. 2009, 5, 2531–2543.

(77) Siuda, I.; Thøgersen, L. Conformational flexibility of the leucine binding protein examined by protein domain coarse-grained molecular dynamics. J. Mol. Model. 2013, 19, 4931–4945.

(78) Humphrey, W.; Dalke, A.; Schulten, K. VMD: Visual molecular dynamics. J. Mol. Graph. 1996, 14, 33–38.

(79) Hsu, P.-C.; Bruininks, B. M. H.; Jefferies, D.; Cesar Telles de Souza, P.; Lee, J.; Patel, D. S.; Marrink, S. J.; Qi, Y.; Khalid, S.; Im, W. CHARMM-GUI Martini Maker for modeling and simulation of complex bacterial membranes with lipopolysaccharides. J. Comput. Chem. 2017, 38, 2354–2363.

(80) Qi, Y.; Ingolfsson, H. I.; Cheng, X.; Lee, J.; Marrink, S. J.; Im, W. CHARMM-GUI Martini Maker for Coarse-Grained Simulations with the Martini Force Field. J. Chem. Theory Comput. 2015, 11, 4486–4494.

(81) Borges-Araujo, L.; Souza, P. C. T.; Fernandes, F.; Melo, M. N. Improved Parameterization of Phosphatidylinositide Lipid Headgroups for the Martini 3 Coarse-Grain Force Field. J. Chem. Theory Comput. 2021, 18, 357–373.

(82) Nose, S. Molecular-Dynamics Method for Simulations in the Canonical Ensemble. Mol. Phys. 1984, 52, 255–268.

(83) Hoover, W. G. Canonical Dynamics - Equilibrium Phase-Space Distributions. Phys. Rev. A 1985, 31, 1695–1697.

(84) Huang, J.; Rauscher, S.; Nawrocki, G.; Ran, T.; Feig, M.; de Groot, B. L.; Grubmüller, H.; MacKerell, A. D., Jr CHARMM36m: an improved force field for folded and intrinsically disordered proteins. Nat. Methods 2017, 14, 71–73.

(85) Hahn, C. DuIvyTools, version 0.3.0. 2022; 10.5281/zenodo.6340263.

(86) Corradi, V.; Mendez-Villuendas, E.; Ingolfsson, H. I.; Gu, R.-X.; Siuda, I.; Melo, M. N.; Moussatova, A.; DeGagne, L. J.; Sejdiu, B. I.; Singh, G.; Wassenaar, T. A.; Delgado Magnero, K.; Marrink, S. J.; Tieleman, D. P. Lipid–Protein Interactions Are Unique Fingerprints for Membrane Proteins. ACS Cent. Sci. 2018, 4, 709–717, PMID:29974066.

(87) Wassenaar, T. A.; Pluhackova, K.; Bockmann, R. A.; Marrink, S. J.; Tieleman, D. P. Going Backward: A Flexible Geometric Approach to Reverse Transformation from Coarse Grained to Atomistic Models. J. Chem. Theory Comput. 2014, 10, 676–690.

(88) Wassenaar, T. A.; Pluhackova, K.; Moussatova, A.; Sengupta, D.; Marrink, S. J.; Tieleman, D. P.; Böckmann, R. A. High-Throughput Simulations of Dimer and Trimer Assembly of Membrane Proteins. The DAFT Approach. J. Chem. Theory Comput. 2015, 11, 2278–2291.

