## Supplementary Information for "Lipopolysaccharide chain length and coat assembly keep guanylate-binding proteins upright on bacterial membranes"

### Supplementary Tables

Table S1: **Summary of the simulations of the closed mGBP2 monomer and dimer at membranes.** For each run, the time interval of the binding event is given together with the domains involved. The number of clusters reflects the number of distinct binding events. MiD/ED TR: MiD/ED transition region.

| | | | |
|---|---|---|---|
| **75%/20%/5% DOPC/CHOL/CL membrane and mGBP2 monomer** | | | |
| System | Binding events [$\mu$s] | Interacting domains | Clusters during the binding events |
| Conf 1, Run 1 | 0.94–10 | CGN insertion (BP1) | 50 |
| Conf 1, Run 2 | 2.80–5.30, 5.31–6.23, 6.24–10 | MiD/ED TR → GD → CGN insertion | 13 & 7 & 26 |
| Conf 1, Run 3 | 3–10 | CGN insertion | 26 |
| Conf 2, Run 1 | 1.1–10 | MiD/ED TR | 58 |
| Conf 2, Run 2 | 1.85–10 | MiD/ED TR | 50 |
| Conf 2, Run 3 | 0.4–10 | CGN insertion | 47 |
| Conf 3, Run 1 | 0.2–7.61, 8.38–10 | MiD/ED TR → Dissociation → MiD/ED TR | 103 & 20 |
| Conf 3, Run 2 | 1.2–6.35, 7.75–10 | MiD/ED TR/GD → Dissociation → GD | 28 & 9 |
| Conf 3, Run 3 | 0.16–10 | CGN insertion | 41 |
| **75%/20%/5% DOPC/CHOL/POPS membrane and mGBP2 monomer** | | | |
| System | Binding events [$\mu$s] | Interacting domains | Clusters during the binding events |
| Conf 1, Run 1 | 0.08–10 | CGN insertion | 49 |
| Conf 1, Run 2 | 8.89–10 | MiD/ED TR | 10 |
| Conf 1, Run 3 | 0.28–10 | CGN insertion | 99 |
| Conf 2, Run 1 | 0.76–10 | CGN insertion | 69 |
| Conf 2, Run 2 | 0.28–10 | CGN insertion | 46 |
| Conf 2, Run 3 | 8.76–10 | CGN insertion | 16 |
| Conf 3, Run 1 | 3.84–10 | MiD/ED TR | 33 |
| Conf 3, Run 2 | 0.17–10 | CGN insertion | 94 |
| Conf 3, Run 3 | – | no interactions | – |
| **80%/20% DOPC/CHOL membrane and mGBP2 monomer** | | | |
| System | Binding events [$\mu$s] | Interacting domains | Clusters during the binding events |
| Conf 1, Run 1 | 9.64–10 | lying with GD on the MEM, no insertion | 7 |
| Conf 1, Run 2 | 0.05–10 | CGN insertion | 63 |
| Conf 1, Run 3 | 1.69–10 | CGN insertion | 77 |
| Conf 2, Run 1 | 5.18–10 | MiD/ED TR | 37 |
| Conf 2, Run 2 | 0.87–1.15, 1.37–10 | MiD/ED TR → Dissociation → CGN insertion | 5 & 50 |
| Conf 2, Run 3 | 1.67–10 | CGN insertion | 22 |
| Conf 3, Run 1 | 3.21–10 | CGN insertion | 41 |
| Conf 3, Run 2 | 1.21–10 | CGN insertion | 89 |
| Conf 3, Run 3 | 1.18–10 | CGN insertion | 55 |
| **75%/20%/5% DOPC/CHOL/CL membrane and mGBP2 dimer** | | | |
| System | Binding events [$\mu$s] | Interacting domains | Clusters during the binding events |
| Conf 1, Run 1 | M1: 4.44–10 | M1: MiD/ED TR | 118 |
| | M2: 5.96–6.33 | M2: Lying on the MEM → Dissociation | |
| Conf 2, Run 1 | M1: 5.37–5.74, 7.50–10 | M1: MiD/ED TR → Dissociation → MiD/ED TR | 145 |
| | M2: 4.56–10 | M2: MiD/ED TR → CGN insertion | |
| Conf 3, Run 1 | M1: – | M1: No association | 135 |
| | M2: 7.00-10 | M2: MiD/ED TR | |

Table S2: **Lipid interactions of the closed mGBP2 monomer.** The binding poses (BP) are BP1: insertion of the geranylgeranyl anchor (CGN); BP2: binding via the MiD/ED transition region (TR); BP3: binding via the GD. The total time spent in each pose is given in brackets. For each run, the total number of interacting key residues is listed, together with the five residues with the longest association times for CL, POPS, or DOPC. The lipid numbers refer to the whole protein during the longest association event on the membrane. Where more than one binding pose occurred, BP1 was used.

| | | | | |
|---|---|---|---|---|
| **75%/20%/5% DOPC/CHOL/CL membrane and mGBP2 monomer** | | | | |
| System | Binding pose (total time [$\mu s$]) | Total Residues (Cutoff: 10%) | Five top residues | Lipid number (whole) |
| Conf 1, Run 1 | BP1 (9.1) | DOPC: 35; CHOL: 22 ; CL: 26 | CGN, K560, K192, K585, K578, K568 | DOPC: 13.3 ± 2.8 CHOL: 3.6 ± 1.5 CL: 7.4 ± 2.5 |
| Conf 1, Run 2 | BP2 (2.5) → BP3 (0.9) → BP1 (3.8) | DOPC: 28; CHOL: 11 ; CL: 9 | CGN, K568, K560, R572, K581, K427 | DOPC: 5.8 ± 3.8 CHOL: 0.8 ± 0.8 CL: 1.9 ± 1.5 (BP2) |
| | | | | DOPC: 12.1 ± 4.0 CHOL: 2.8 ± 1.6 CL: 5.6 ± 2.5 (BP1) |
| Conf 1, Run 3 | BP1 (7.0) | DOPC: 34; CHOL: 18 ; CL: 24 | CGN, K585, R553, K560, K568, R231 | DOPC: 11.8 ± 5.2 CHOL: 3.1 ± 2.0 CL: 6.0 ± 3.4 |
| Conf 2, Run 1 | BP2 (8.9) | DOPC: 31; CHOL: 9 ; CL: 5 | K427, T424, P428, S426, Q478 | DOPC: 7.4 ± 3.8 CHOL: 1.0 ± 1.0 CL: 2.6 ± 1.7 |
| Conf 2, Run 2 | BP2 (8.2) | DOPC: 30; CHOL: 9 ; CL: 7 | K427, T424, P428, K443, S426 | DOPC: 8.1 ± 4.1 CHOL: 1.2 ± 1.1 CL: 2.8 ± 1.8 |
| Conf 2, Run 3 | BP1 (9.6) | DOPC: 37; CHOL: 20 ; CL: 29 | K560, CGN, K192, H202, K585, K578 | DOPC: 12.5 ± 4.7 CHOL: 3.0 ± 1.5 CL: 6.8 ± 2.9 |
| Conf 3, Run 1 | BP2 (9.0) | DOPC: 20; CHOL: 6 ; CL: 5 | K427, K421, L422, S426, T424 | DOPC: 6.4 ± 4.2 CHOL: 0.9 ± 0.9 CL: 2.7 ± 1.9 |
| Conf 3, Run 2 | BP2 (5.1) → BP3 (2.3) | DOPC: 19 CHOL: 3 CL: 4 | K213, K207, K427, F217, L422 | DOPC: 6.7 ± 4.0 CHOL: 0.7 ± 0.8 CL: 2.0 ± 1.6 |
| | | | | DOPC: 6.9 ± 3.6 CHOL: 1.3 ± 0.8 CL: 2.3 ± 1.5 (BP1) |
| Conf 3, Run 3 | BP1 (9.2) | DOPC: 102 CHOL: 45 CL: 46 | CGN, K568, R553, K560, K230, R572 | DOPC: 21.9 ± 8.4 CHOL: 4.5 ± 2.5 CL: 10.1 ± 4.9 |
| **75%/20%/5% DOPC/CHOL/POPS membrane and mGBP2 monomer** | | | | |
| System | Binding pose (total time [$\mu s$]) | Total Residues (Cutoff: 10%) | Five top residues | Lipid number (whole) |
| Conf 1, Run 1 | BP1 (9.9) | DOPC: 74 CHOL: 36 POPS: 7 | CGN, K581, K578, K585, S582, K534 | DOPC: 29.6 ± 6.0 CHOL: 2.1 ± 1.5 POPS: 10.8 ± 3.0 |
| Conf 1, Run 2 | BP2 (1.1) | DOPC: 2 CHOL: – POPS: – | P428, K427 | DOPC: 12.8 ± 2.1 CHOL: 0.4 ± 0.7 POPS: 4.2 ± 1.7 |
| Conf 1, Run 3 | BP1 (9.7) | DOPC: 65 CHOL: 22 POPS: 5 | CGN, K560, K585, K568, R572, L559 | DOPC: 23.7 ± 8.5 CHOL: 1.9 ± 1.4 POPS: 8.7 ± 3.1 |
| Conf 2, Run 1 | BP1 (9.2) | DOPC: 38 CHOL: 17 POPS: 6 | CGN, K585, K568, K560, R572, K208 | DOPC: 14.8 ± 5.0 CHOL: 1.6 ± 1.4 POPS: 5.9 ± 2.6 |
| Conf 2, Run 2 | BP1 (9.7) | DOPC: 40 CHOL: 17 POPS: 5 | CGN, K560, K585, K568, K192, K208 | DOPC: 13.0 ± 5.8 CHOL: 1.3 ± 1.2 POPS: 5.1 ± 2.7 |
| Conf 2, Run 3 | BP1 (1.2) | DOPC: 2 CHOL: – POPS: – | CGN, K585, K568, K560, S582, R572 | DOPC: 11.9 ± 4.1 CHOL: 1.6 ± 1.3 POPS: 4.6 ± 2.4 |
| Conf 3, Run 1 | BP2 (6.1) | DOPC: 18 CHOL: 7 POPS: – | K427, L422, K421, P428, T476 | DOPC: 6.7 ± 4.8 CHOL: 0.5 ± 0.8 POPS: 2.3 ± 1.9 |
| Conf 3, Run 2 | BP1 (9.8) | DOPC: 39 CHOL: 17 POPS: 2 | CGN, K585, R572, K568, K560, K192 | DOPC: 13.5 ± 5.6 CHOL: 1.4 ± 1.2 POPS: 5.1 ± 3.0 |
| Conf 3, Run 3 | no association | – | – | – |
| **80%/20% DOPC/CHOL membrane and mGBP2 monomer** | | | | |
| System | Binding pose (total time [$\mu s$]) | Total Residues (Cutoff: 10%) | Five top residues | Lipid number (whole) |
| Conf 1, Run 1 | lying with GD on the MEM | – | – | – |
| Conf 1, Run 2 | BP1 (9.9) | DOPC: 30 CHOL: 16 | CGN, K585, K568, R572, K560, L559 | DOPC: 14.3 ± 4.6 CHOL: 4.6 ± 2.3 |
| Conf 1, Run 3 | BP1 (8.3) | DOPC: 31 CHOL: 17 | CGN, K585, K568, K560, R572, L559 | DOPC: 17.0 ± 3.8 CHOL: 6.4 ± 2.2 |
| Conf 2, Run 1 | BP2 (4.8) | DOPC: 26 CHOL: 7 | K427, Y432, P428, S479, Q478 | DOPC: 9.4 ± 4.3 CHOL: 2.9 ± 1.8 |
| Conf 2, Run 2 | BP2 (0.2) → BP1 (8.6) | DOPC: 15 CHOL: 4 | CGN, K585, K568, R572, E564, K569 | DOPC: 11.0 ± 5.1 CHOL: 3.4 ± 2.4 (BP1) |
| Conf 2, Run 3 | BP1 (8.3) | DOPC: 31 CHOL: 15 | CGN, K568, R572, K560, L559, K578 | DOPC: 15.8 ± 3.8 CHOL: 5.6 ± 1.9 |
| Conf 3, Run 1 | BP1 (6.8) | DOPC: 31 CHOL: 13 | CGN, K568, K585, R572, K560, I571 | DOPC: 13.4 ± 4.6 CHOL: 4.6 ± 2.5 |
| Conf 3, Run 2 | BP1 (8.8) | DOPC: 31 CHOL: 17 | CGN, K585, K568, K560, R572, L559 | DOPC: 13.9 ± 5.2 CHOL: 5.1 ± 2.6 |
| Conf 3, Run 3 | BP1 (8.8) | DOPC: 38 CHOL: 19 | CGN, K585, K568, K560, R572, K192 | DOPC: 17.3 ± 4.6 CHOL: 6.1 ± 2.6 |

### Supplementary Figures

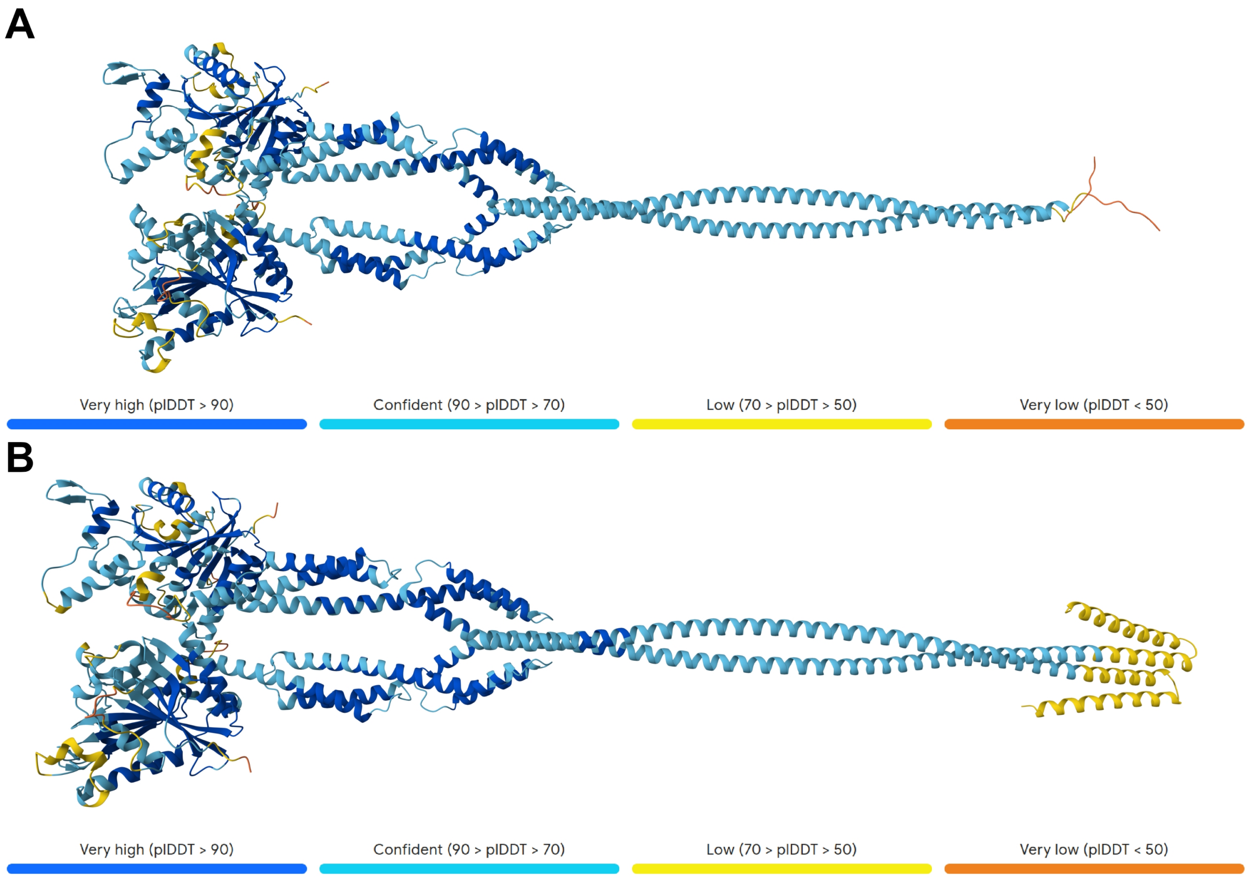


Figure S1: **AlphaFold 3 predictions of mGBP2 and mGBP7 dimers.** AlphaFold 3 predicts the **(A)** mGBP2 dimer and **(B)** mGBP7 dimer with extended EDs but without criss-crossing of the GDs and MiDs of the two protomers.

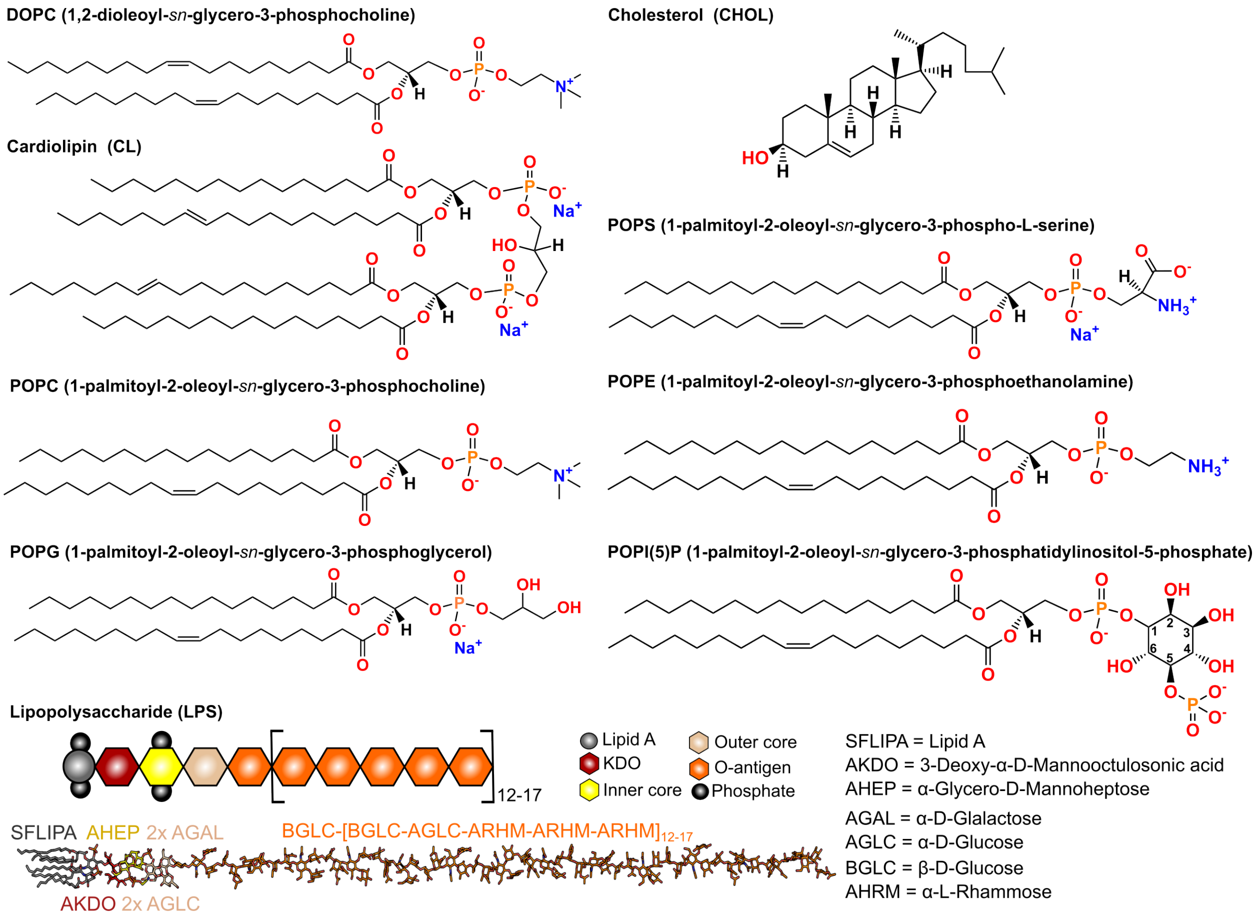


Figure S2: **Chemical identities of the lipids used to construct the membrane systems.**

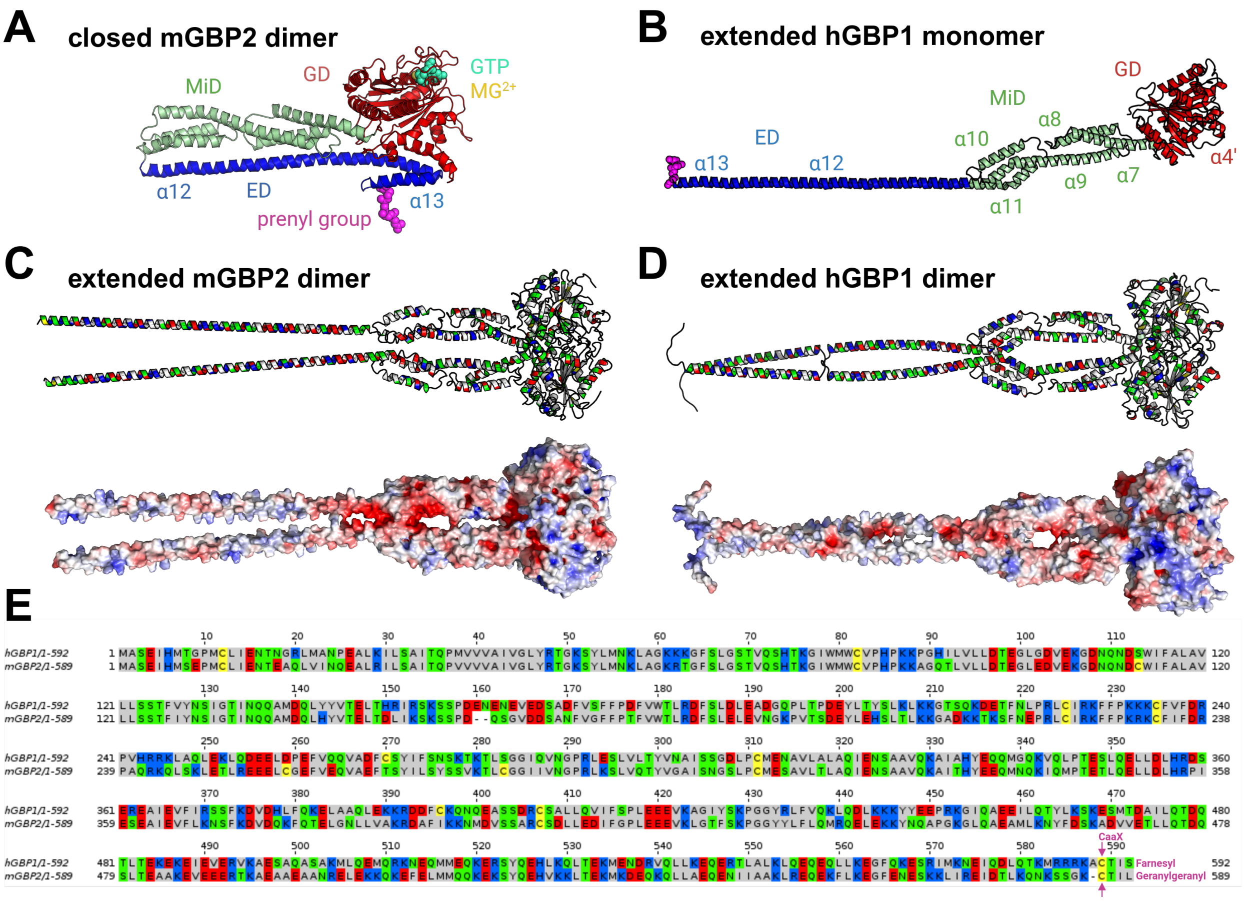


Figure S3: **Structural features and residue analysis of hGBP1 and mGBP2. (A)** mGBP2 in the closed form and **(B)** hGBP1 in the extended form, colored by domain: GD (red), MiD (green), ED (blue), and lipidation (magenta). **(C, D)** Residue-type coloring and electrostatic potential maps of the extended dimeric hGBP1 (C) and mGBP2 (D). Amino acids are colored green for polar, yellow for cysteine, red for negatively charged, blue for positively charged, and gray for neutral residues. The electrostatic potential was calculated with the PyMOL plugin tool[1,2] and is shown between −5 (red) and +5 kT/e (blue) for all proteins. **(E)** Sequences of hGBP1 (top) and mGBP2 (bottom), using the same residue coloring as in (C, D) and magenta for the prenylation site. The two proteins share a sequence identity of 408/599 (68.11%) and a sequence similarity of 489/599 (81.64%).

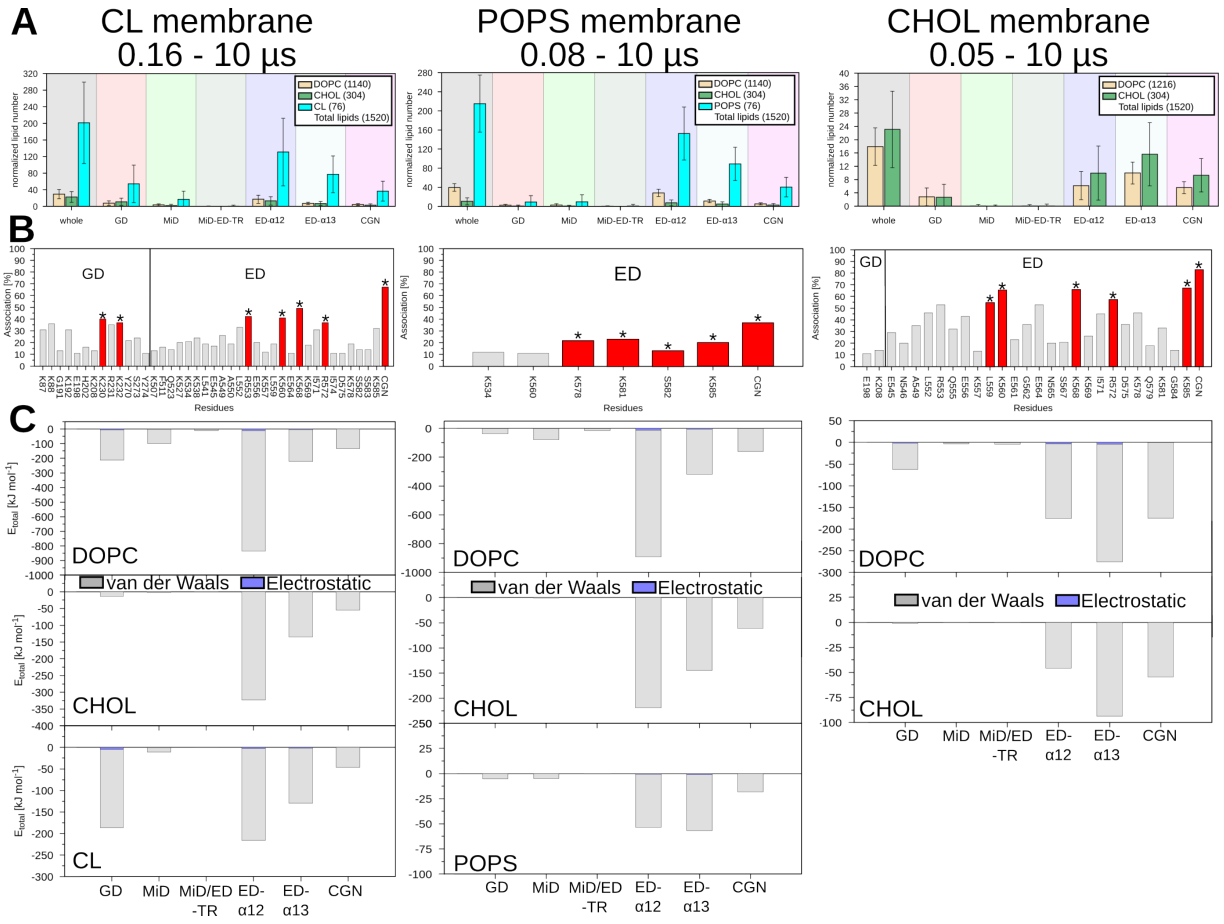


Figure S4: **Lipid numbers, association times, and interaction energies for the closed mGBP2 monomer.** **(A)** Normalized number of each lipid over time for each domain and for the whole protein, with error bars, for one representative run per membrane in which mGBP2 binds via the geranylgeranyl (CGN) anchor. The total lipid numbers are given in the legend. **(B)** Association time of the residues, in percent, for the same runs, with a cut-off of 10%. Red boxes marked with an asterisk indicate the five or six key residues plus the geranylgeranyl anchor with the highest association times; residues are assigned to their domain. **(C)** Interaction energies of each lipid for the different domains and helices, with van der Waals contributions in gray and electrostatic contributions in blue. The corresponding values for the other runs are summarized in **Table S2**.

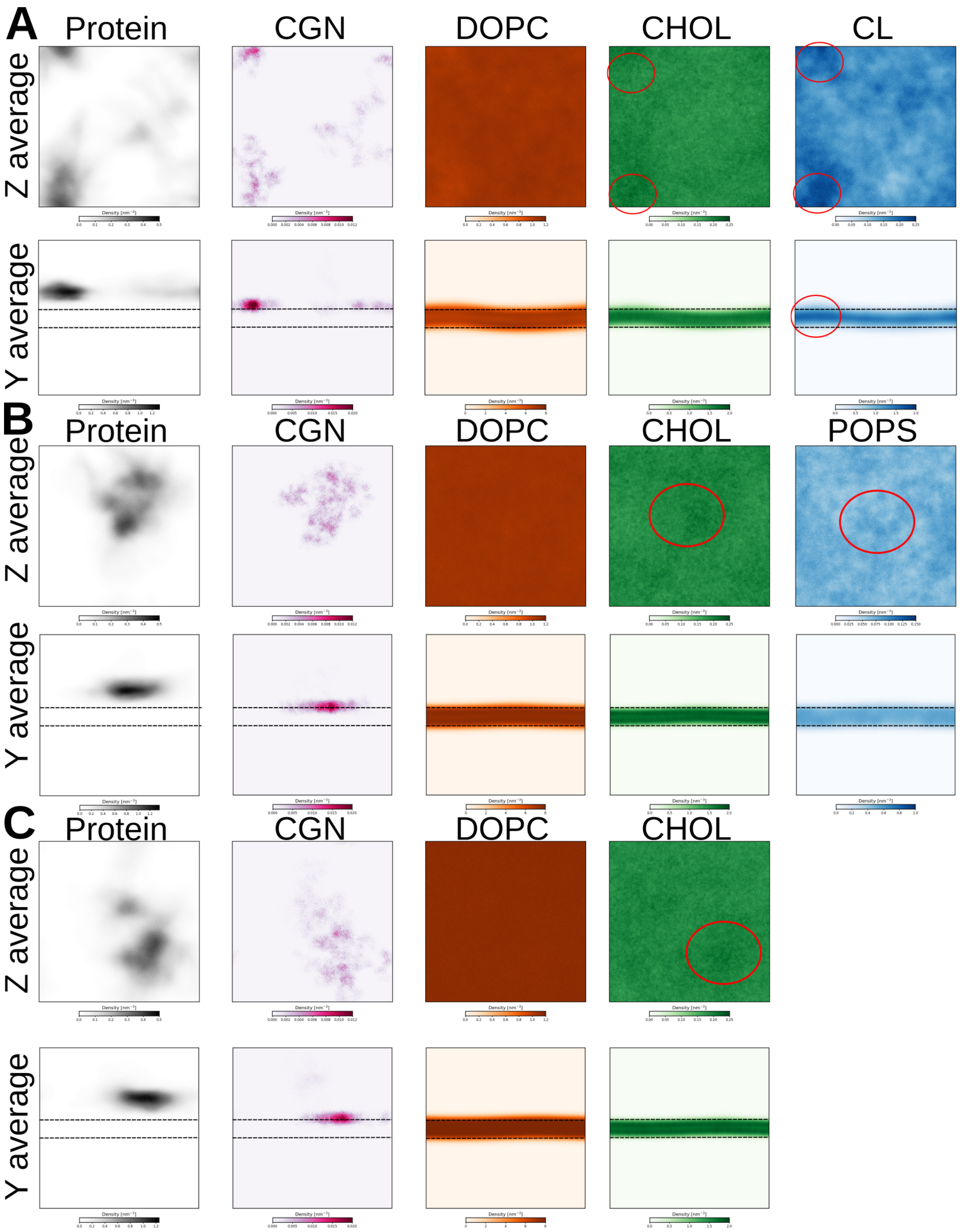


Figure S5: **Density maps for the closed mGBP2 monomer on the three membranes.** The Z average gives the number of molecules per $nm^3$ in top view, showing the clustering of protein and lipids, while the Y average gives the side view of the membrane. For one representative run of **(A)** CL-mem, **(B)** PS-mem, and **(C)** PC-mem, the two rows show the Z and Y averages, while the columns represent protein and its geranylgeranyl anchor (CGN), followed by the individual lipids. The red circles mark the lipid areas overlapping with mGBP2. In the side views, membrane undulations are visible beneath the bound protein, most pronounced on CL-mem; the corresponding protein-free reference simulations are shown in **Fig. S6**.

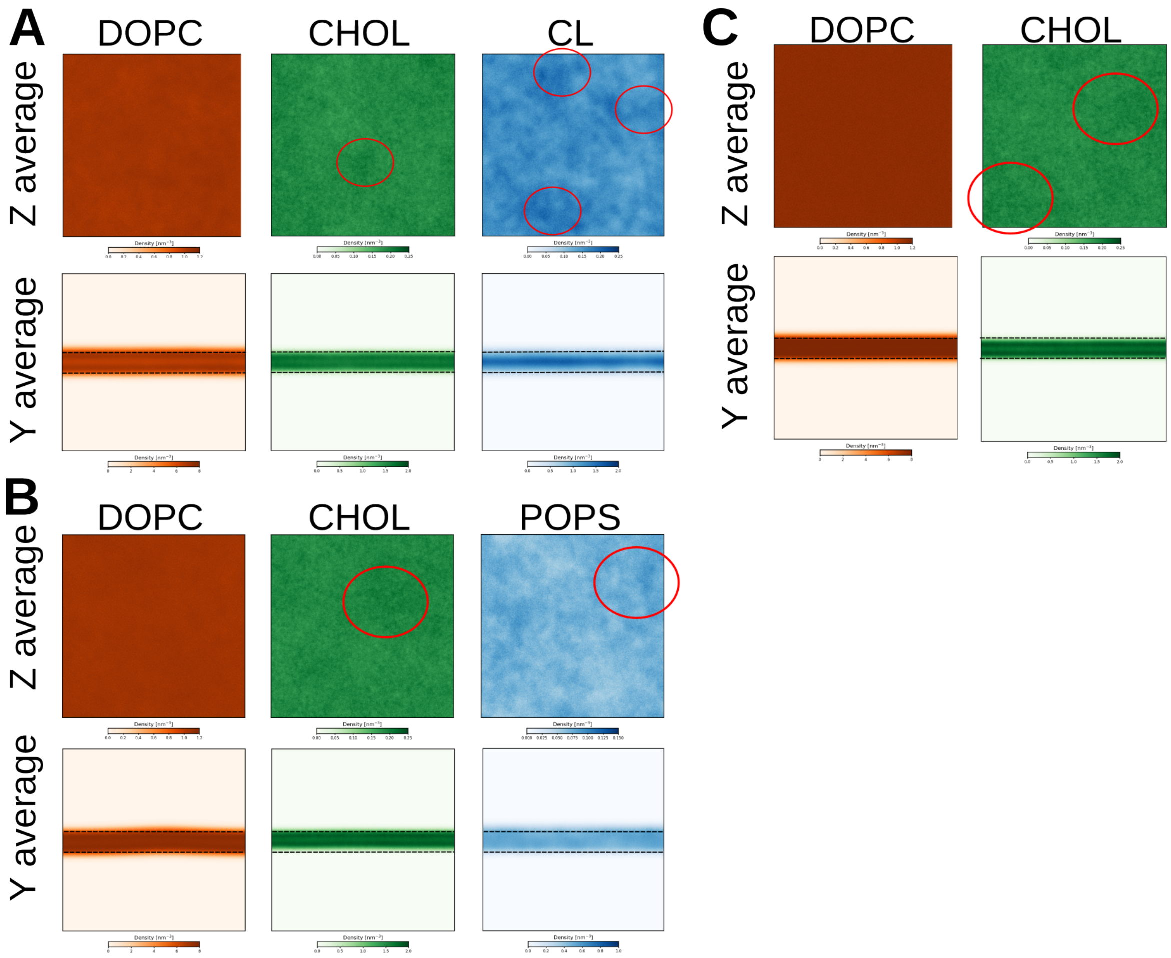


Figure S6: **Density maps for the protein-free reference membranes.** Density maps as in **Fig. S5**, for the three membranes simulated without mGBP2: **(A)** CL-mem, **(B)** PS-mem, and **(C)** PC-mem. The membranes remain flat and show only weak lipid clustering (red circles), in contrast to the simulations with mGBP2.

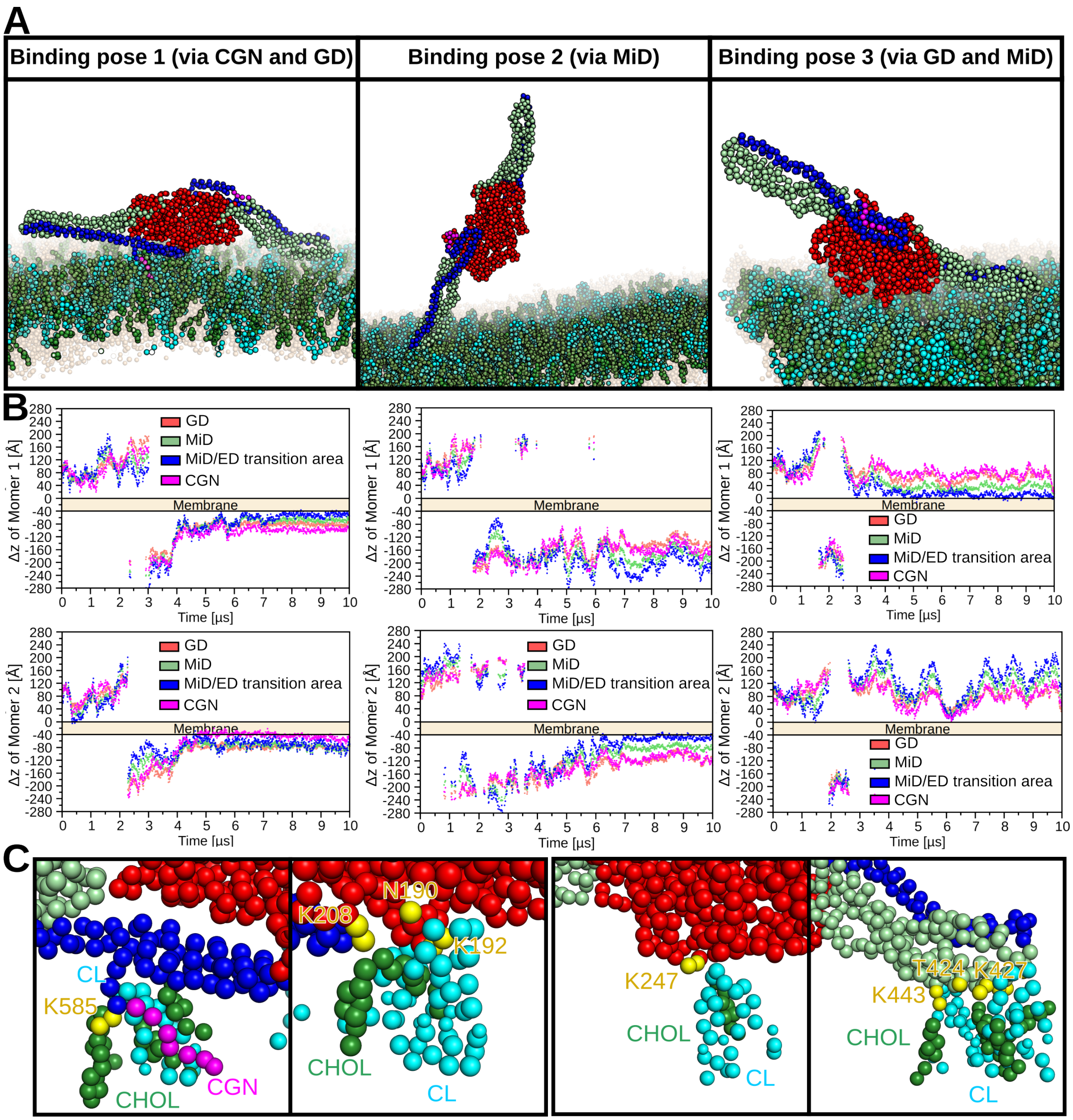


Figure S7: **Interactions of the closed mGBP2 dimer on CL-mem. (A)** The dimer adopts the same binding poses (BP1–BP3) as the monomer. **(B)** Time evolution of the Δz-position for BP1, BP2, and BP3. CGN denotes the geranylgeranyl anchor, and the membrane is indicated as a shaded band. **(C)** View of the interactions with CL and CHOL.

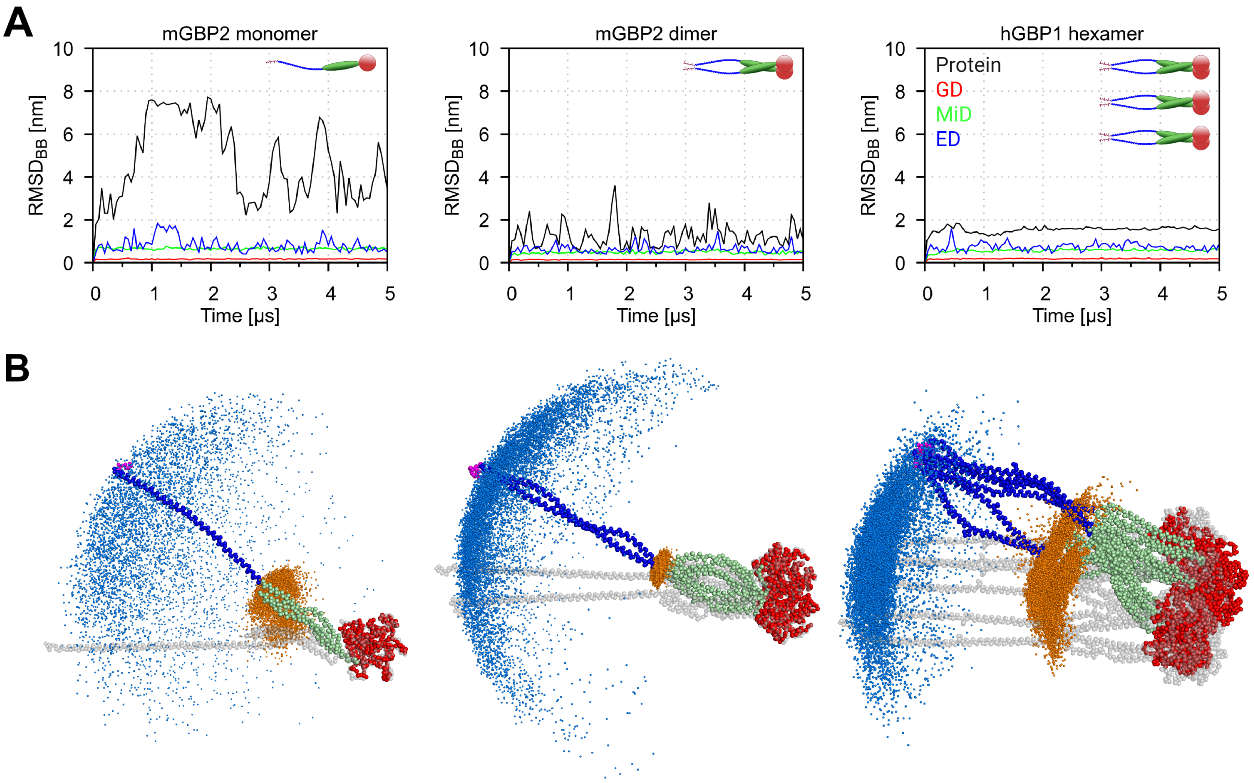


Figure S8: **Stability of the open mGBP2 and hGBP1 structures in solution. (A)** Time evolution of the RMSD for the open mGBP2 monomer and dimer and the hGBP1 hexamer in solution. The RMSD was computed for the entire protein (black), the GD (red), the MiD (green), and the ED (blue), in each case after alignment on the corresponding domain, using the backbone beads. An inset cartoon illustrates the protein parts under investigation. **(B)** Spatial distribution of residues 480 and 582 of all protein chains during the CGMD simulations, shown in orange for residue 480 and in marine for residue 582.

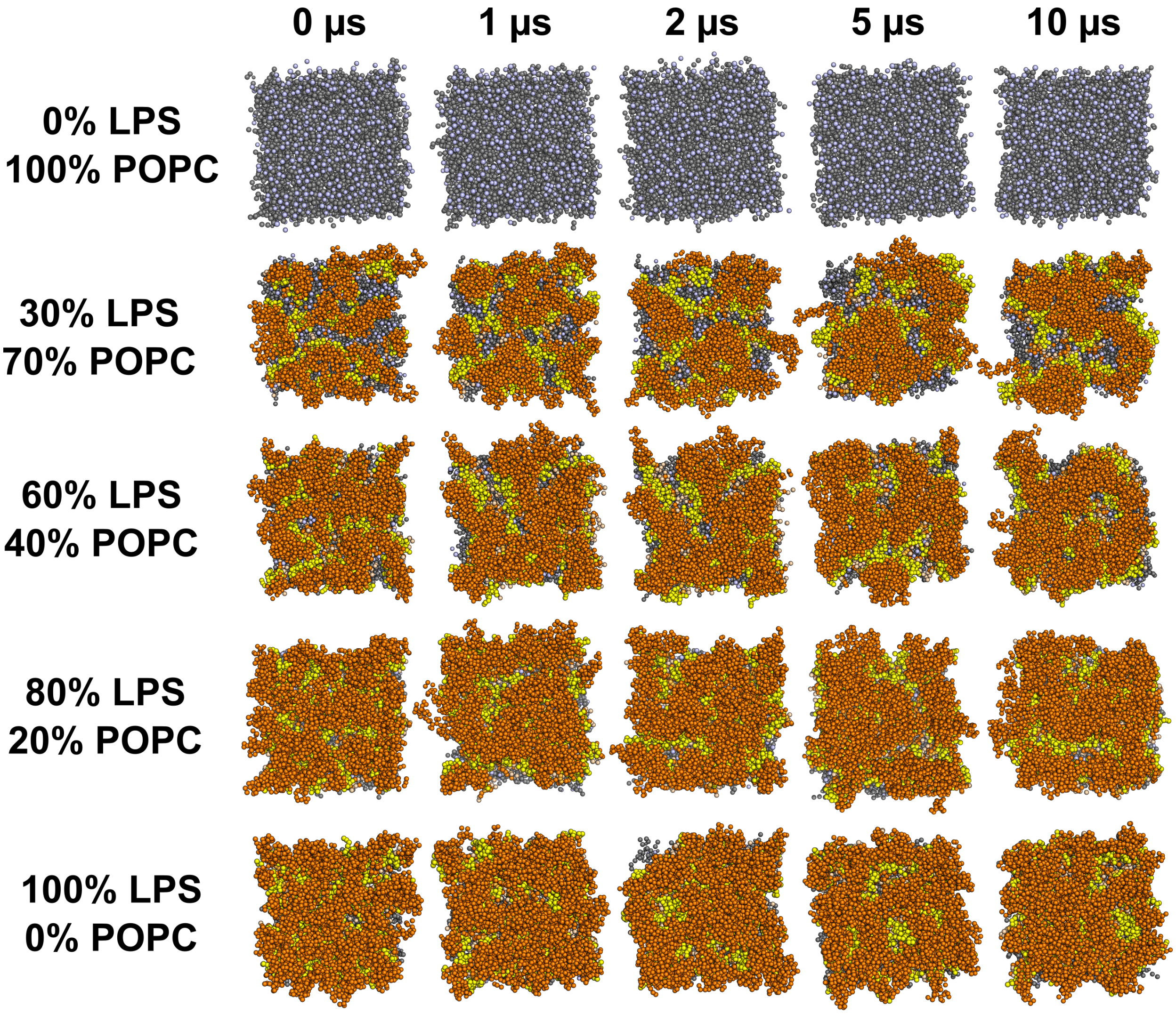


Figure S9: **Protein-free LPS titration simulations.** Top view of the outer leaflet of a pure symmetric POPC bilayer and asymmetric bilayers with 30%, 60%, 80%, and 100% LPS (and 70%, 40%, 20%, and 0% POPC) in the outer leaflet. The inner leaflet consisted of 100% POPC. Each system was simulated for 10 $\mu$s; snapshots at 0, 1, 2, 5, and 10 $\mu$s are shown. The 80% LPS composition, which yielded a well-covered outer leaflet, was used for the LPS membranes in the protein simulations, with POPE instead of POPC as the phospholipid component.

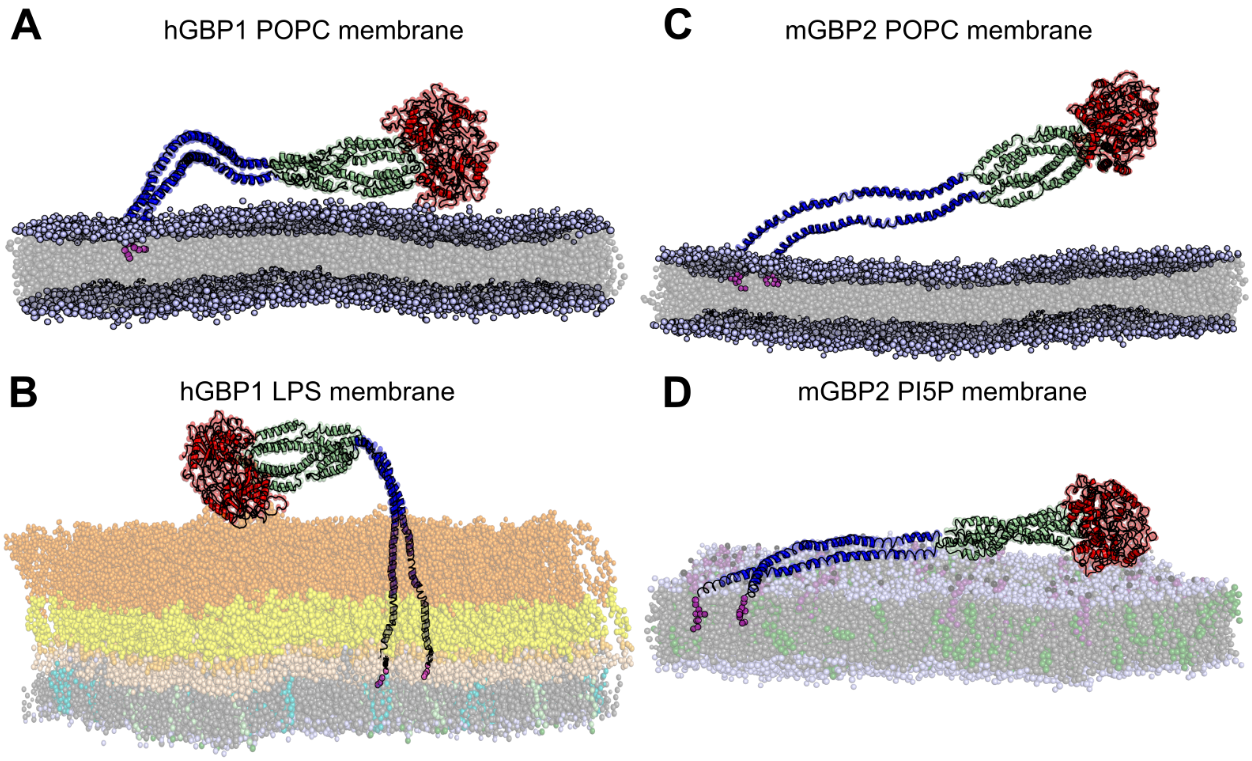


Figure S10: **Backmapped representative snapshots of the open hGBP1 and mGBP2 dimers at membranes.** **(A)** The hGBP1 dimer on the POPC membrane, **(B)** the hGBP1 dimer on the LPS membrane, **(C)** the mGBP2 dimer on the POPC membrane, and **(D)** the mGBP2 dimer on the PI5P membrane. We used simple POPC membranes as reference. The LPS membrane is composed of 80% LPS with short O-antigen chains and 20% POPE in the upper leaflet and 90% POPE, 5% POPG, and 5% CL in the lower leaflet. The PI5P membrane is composed of 5% PI5P, 20% CHOL, and 75% DOPC in the upper leaflet and 20% CHOL and 80% DOPC in the lower leaflet. The corresponding snapshots of the monomers are shown in **Fig. 4A**.

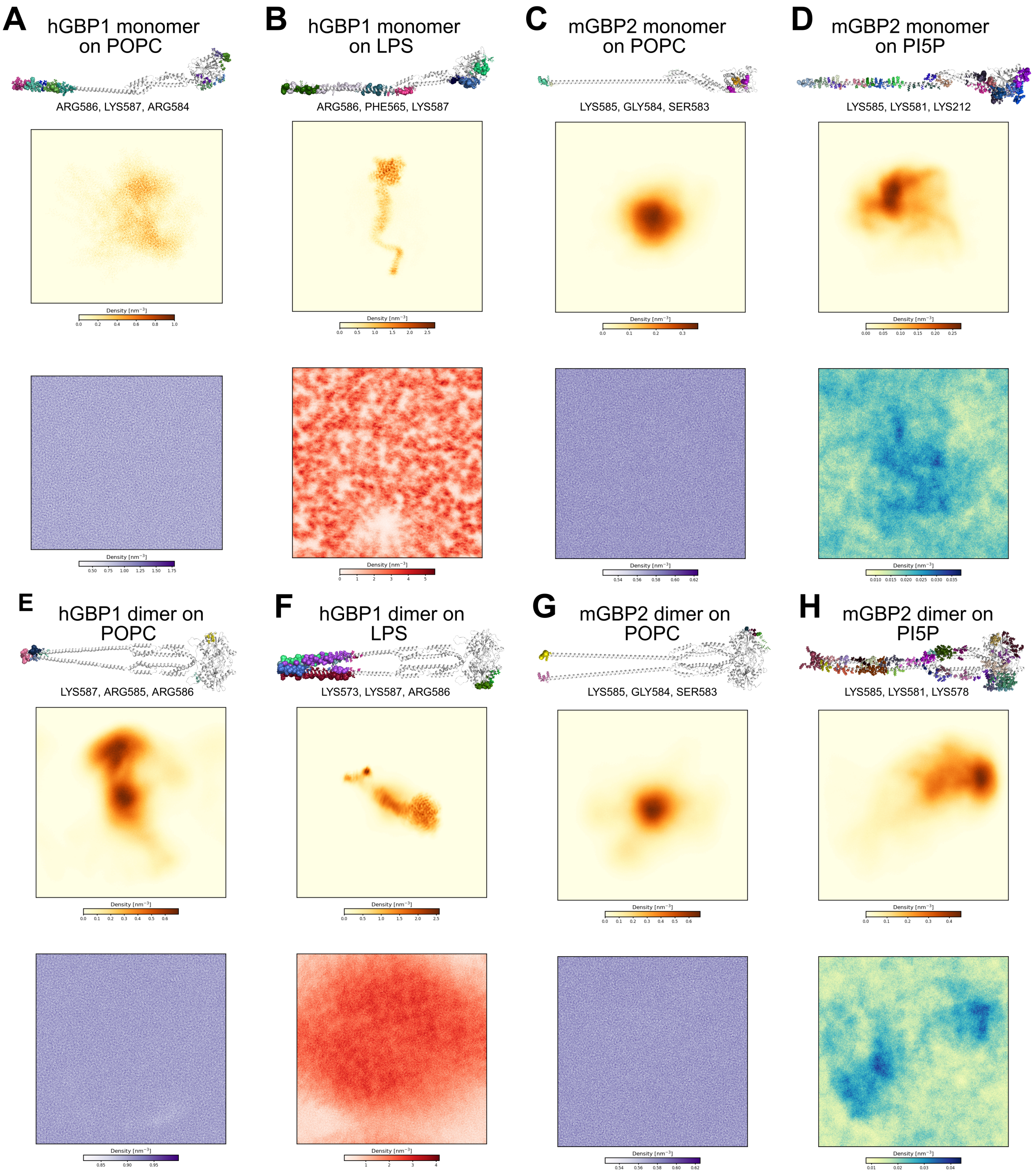


Figure S11: **Binding sites and density maps for the open GBP monomers and dimers at membranes.** Protein–membrane binding sites with different colors, and the three residues of highest lipid occupancy indicated, and density maps of the simulation components (protein and lipid of interest) for **(A)** the hGBP1 monomer on the POPC membrane, **(B)** the hGBP1 monomer on the LPS membrane with short O-antigen chains, **(C)** the mGBP2 monomer on the POPC membrane, **(D)** the mGBP2 monomer on the PI5P membrane, **(E)** the hGBP1 dimer on the POPC membrane, **(F)** the hGBP1 dimer on the LPS membrane, **(G)** the mGBP2 dimer on the POPC membrane, and **(H)** the mGBP2 dimer on the PI5P membrane.

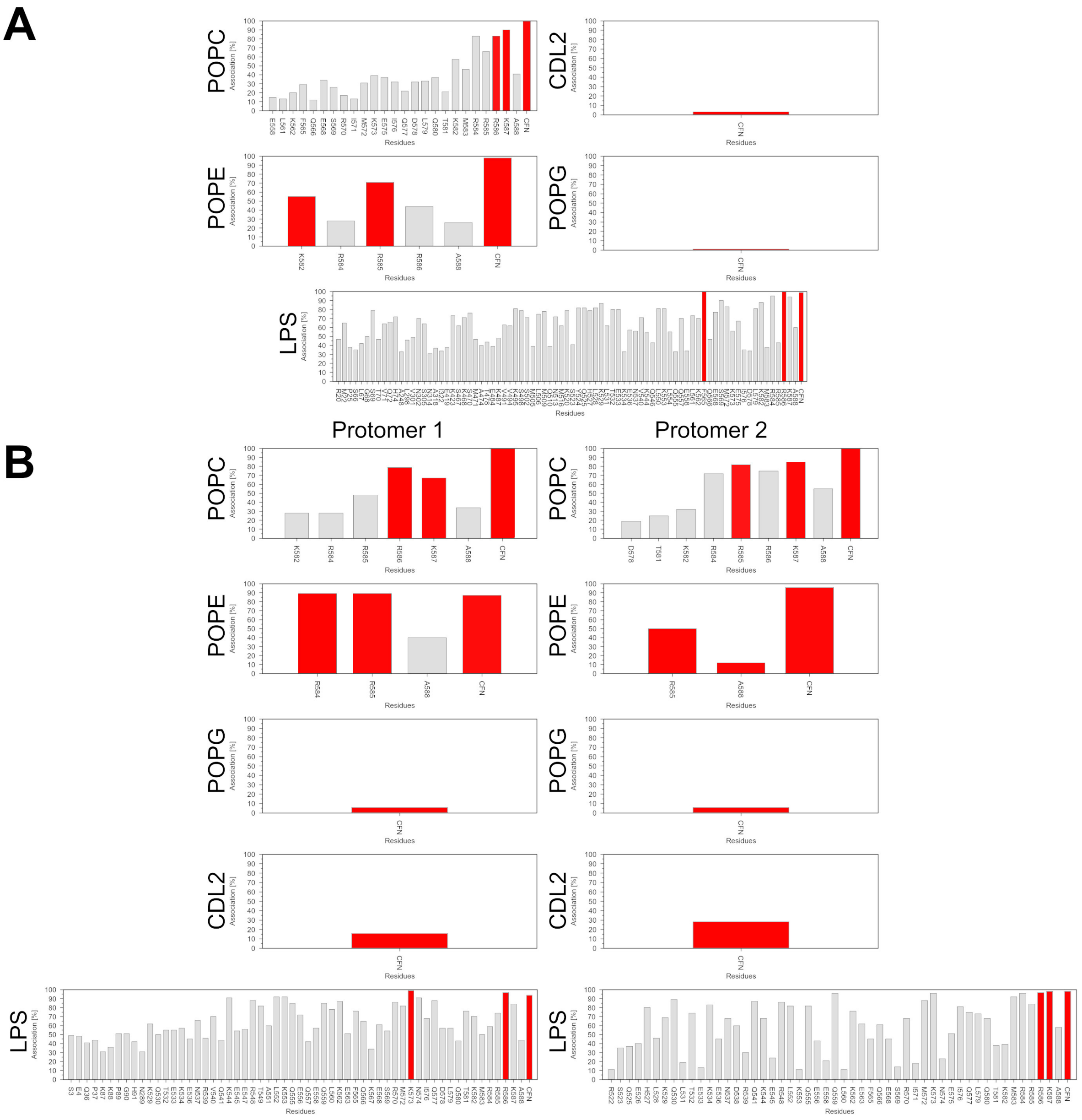


Figure S12: **Association times of the open hGBP1 monomer and dimer on the POPC and LPS membranes.** **(A)** The hGBP1 monomer and **(B)** the hGBP1 dimer on both membranes. The association time is given in percent of the simulation time; only residues with an association time of at least 10% are shown, and the three residues with the highest values are indicated in red. The POPC membrane serves as reference; the asymmetric LPS membrane carries short O-antigen chains (4 repeat units) and is composed of 80% LPS and 20% POPE in the upper leaflet and 90% POPE, 5% POPG, and 5% CL in the lower leaflet.

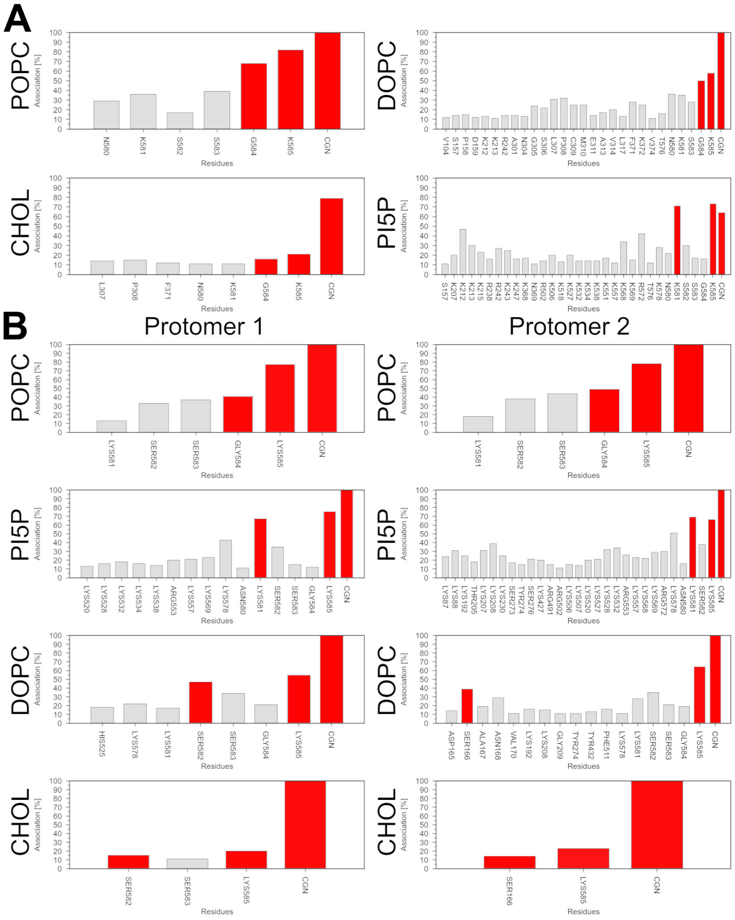


Figure S13: **Association times of the open mGBP2 monomer and dimer on the POPC and PI5P membranes. (A)** The mGBP2 monomer and **(B)** the mGBP2 dimer on both membranes. The association time is given in percent of the simulation time; only residues with an association time of at least 10% are shown, and the three residues with the highest values are indicated in red. The POPC membrane serves as reference; the asymmetric PI5P membrane is composed of 5% PI5P, 20% CHOL, and 75% DOPC in the upper leaflet and 20% CHOL and 80% DOPC in the lower leaflet.

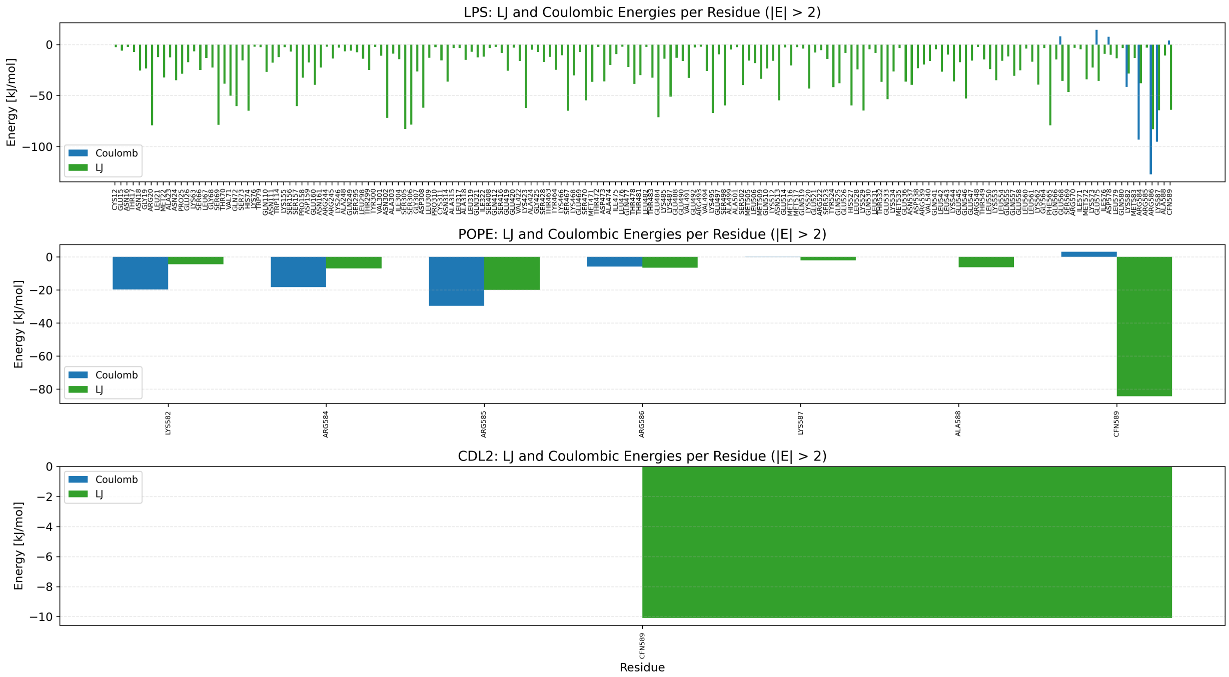


Figure S14: **Interaction energies per lipid and per residue for the open hGBP1 monomer on the LPS membrane.** The energies are plotted with their Coulomb and Lennard-Jones contributions in blue and green, respectively. A cut-off of $|E| > 2$ kJ mol$^{-1}$ was applied to show only significant interactions; for POPG, no contributions within this range were found. CFN589 denotes the farnesyl anchor attached to C589. The width of the bars carries no meaning.

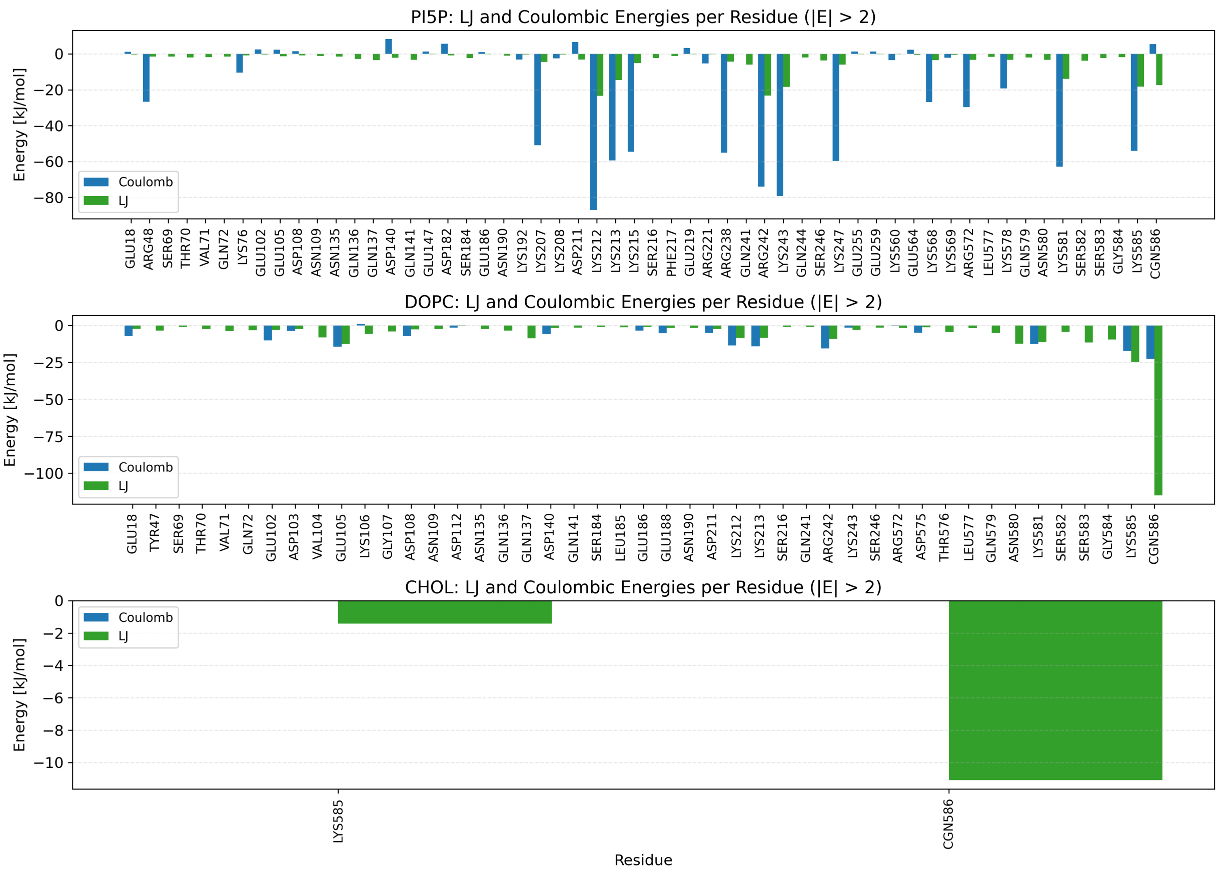


Figure S15: **Interaction energies per lipid and per residue for the open mGBP2 monomer on the PI5P membrane.** The energies are plotted with their Coulomb and Lennard-Jones contributions in blue and green, respectively. A cut-off of $|E| > 2$ kJ mol$^{-1}$ was applied to show only significant interactions. CGN586 denotes the geranylgeranyl anchor attached to C586. The width of the bars carries no meaning.

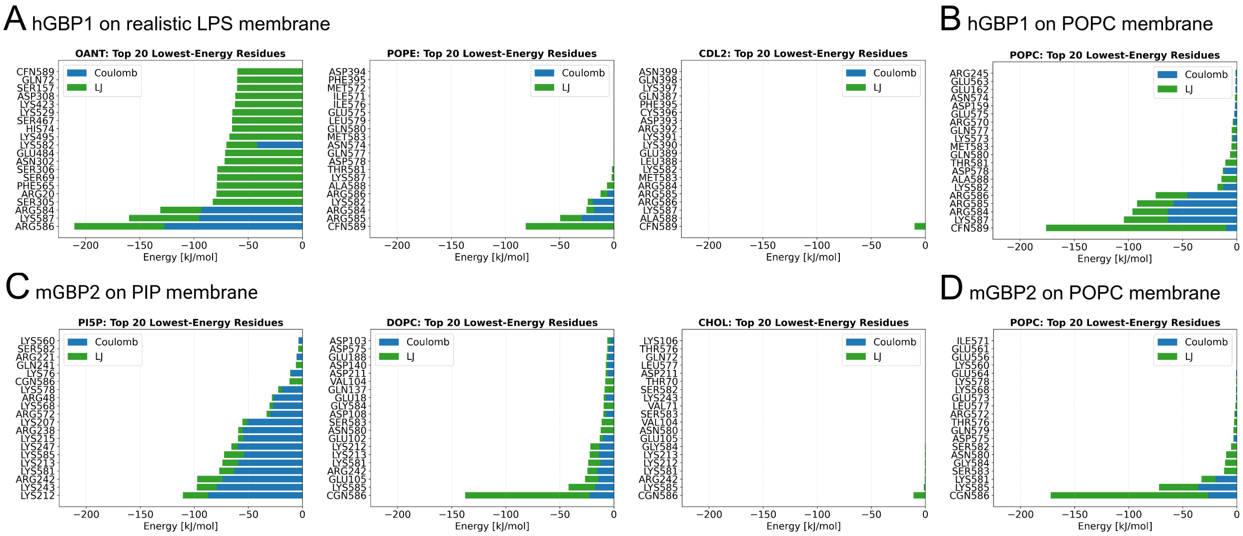


Figure S16: **Top interacting residues of the open hGBP1 and mGBP2 monomers on the different membranes.** The 20 residues with the strongest interaction energies are shown for **(A)** hGBP1 on the LPS membrane, **(B)** hGBP1 on the POPC membrane, **(C)** mGBP2 on the PI5P membrane, and **(D)** mGBP2 on the POPC membrane, resolved by membrane component.

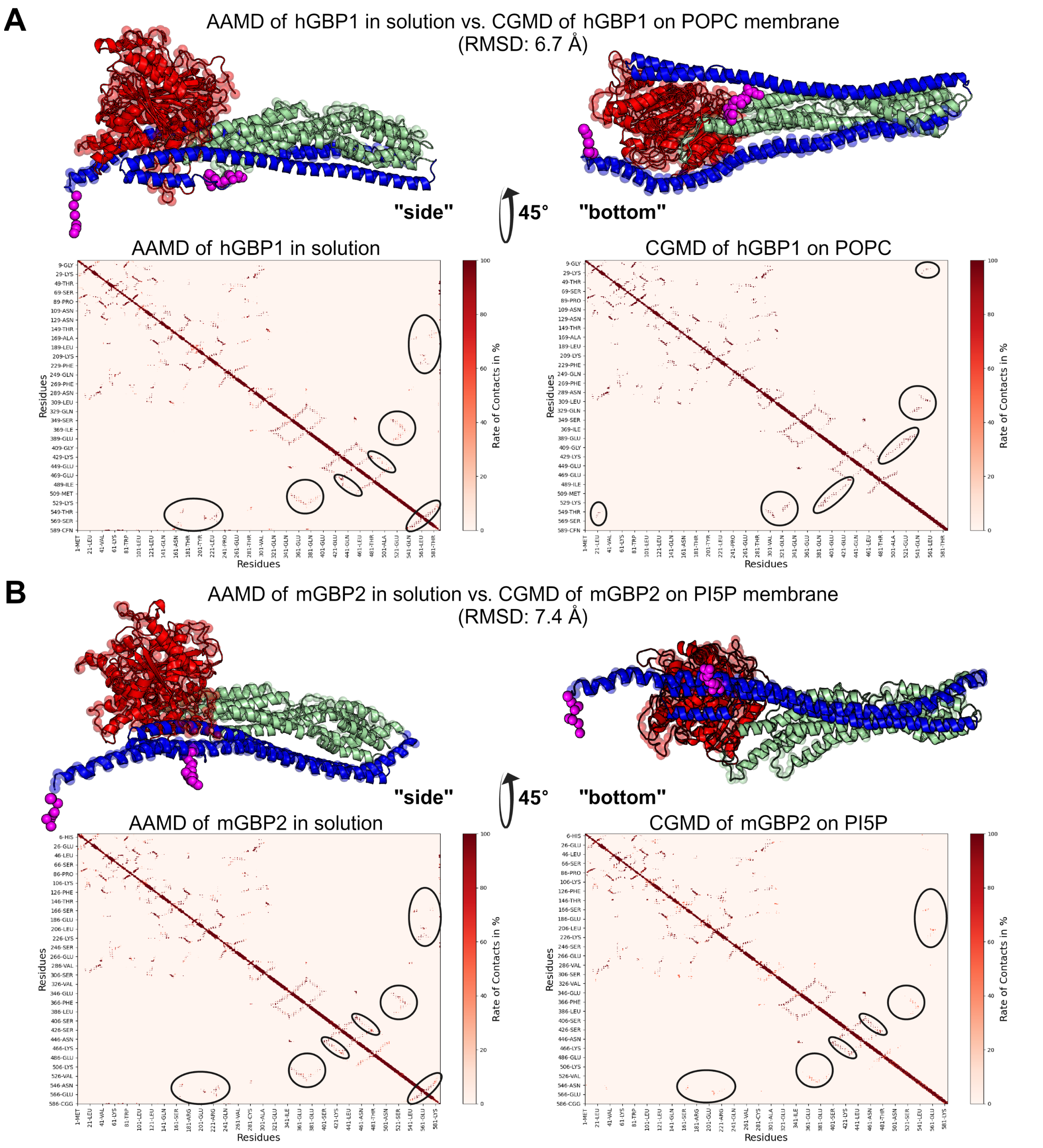


Figure S17: **Closed states of hGBP1 and mGBP2 formed at membranes.** **(A)** Overlay of the closed state of hGBP1 formed after fold-back on the POPC membrane (CGMD, backmapped) with the AAMD structure from our previous work,[3] which retains the ED orientation of the crystal structure (PDB: 1DG3), shown in side and bottom view. In the state formed here, the ED occupies the side of the GD and MiD opposite to that of the crystal structure, and $\alpha$12 and $\alpha$13 form one continuous helix. Intramolecular contact maps of both structures are shown below. **(B)** Overlay of the closed state of mGBP2 formed on the PI5P membrane with the AAMD structure from our previous work,[4] in which the ED retains its crystal-structure orientation, with the corresponding contact maps. GBP domains are colored as in **Fig. 1**.

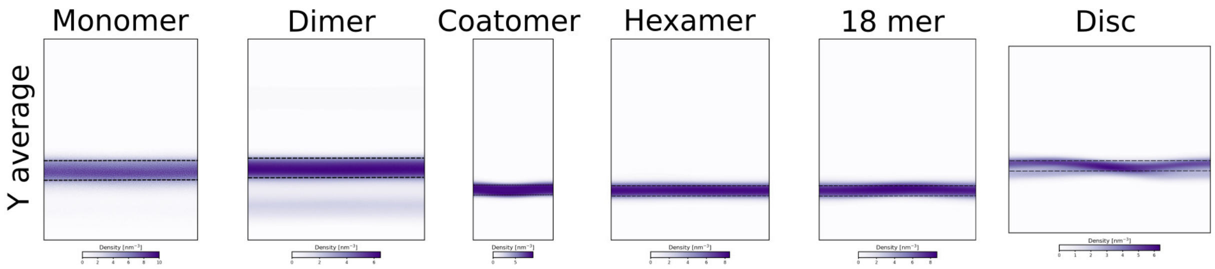


Figure S18: **Density maps of the POPC membrane in the simulations with hGBP1.** Displayed is the Y average density of the POPC lipids in the simulations with the hGBP1 monomer, dimer, hexamer, octadecamer, coat-like assembly, and disc.

### References


(1) Jurrus, E. et al. Improvements to the APBS biomolecular solvation software suite. *Protein Sci.* **2017**, *27*, 112–128.

(2) Dolinsky, T. J.; Nielsen, J. E.; McCammon, J. A.; Baker, N. A. PDB2PQR: an automated pipeline for the setup of Poisson-Boltzmann electrostatics calculations. *Nucleic Acids Res.* **2004**, *32*, W665–W667.

(3) Barz, B.; Loschwitz, J.; Strodel, B. Large-scale, dynamin-like motions of the human guanylate binding protein 1 revealed by multi-resolution simulations. *PLoS Comput. Biol.* **2019**, *15*, 1–29.

(4) Loschwitz, J.; Steffens, N.; Wang, X.; Schäffler, M.; Pfeffer, K.; Degrandi, D.; Strodel, B. Domain motions, dimerization, and membrane interactions of the murine guanylate binding protein 2. *Sci. Rep.* **2023**, *13*, 1–20.
